# SARM1 Activation Promotes Axonal Degeneration Via a Two-Step Liquid-to-Solid Phase Transition

**DOI:** 10.1101/2024.11.19.624255

**Authors:** Zhang Wenbin, Zhou Qinyi, Zhang Jun, Wang Jiachen, Zheng Sanduo, Wang Xiaodong

**Affiliations:** National Institute of Biological Sciences, Beijing; National institute of Biological Sciences, Beijing; National Institute of Biological Sciences

## Abstract

SARM1 protein plays a central role in axonal degeneration, a key process in many neurodegenerative diseases and nerve injuries. It mediates this by depleting axonal NAD+ through its NADase activity, catalyzed by the TIR domain. Normally, this activity is kept in an inactive state and becomes activated in response to various neuronal damage signals. However, the molecular mechanism behind SARM1 activation, particularly how activation is restricted to damaged axons, remains to be fully elucidated. In this study, using a class of pyridine-containing compounds that induce SARM1-dependent cell death and axonal degeneration, we reveal a two-step process of SARM1 activation. The first step involves sub-lethal “priming” activation of SARM1 mediated by the NAD+ precursor NMN, leading to the formation of covalent conjugates between the hydrolyzed product of NAD^+^, adenosine diphosphate ribose (ADPR), and the compounds catalyzed by the intrinsic base exchange activity of SARM1. In the second step, these ADPR-conjugates act as molecular glues, promoting the formation of self-proliferating superhelical filaments. On these filaments, SARM1’s TIR domains adopt an active NADase configuration. As the superhelical filaments rapidly reach their limit of solubility, they precipitate out of the liquid phase as condensates with stable, fully activated NAD hydrolysis activity. Interestingly, we found that a series of reported SARM1 inhibitors currently under clinical development, which target the TIR enzymatic domain, can paradoxically activate SARM1’s NADase activity via this mechanism. These findings provide new insights into how SARM1 activation is spatially restricted to damaged axons and offer important implications for the development of therapeutics targeting SARM1.

## Main

SARM1 (Sterile Alpha and TIR Motif Containing 1) protein is known as a key mediator of axon degeneration during nerve injury^1,2^. It comprises an auto-inhibitory N-terminal armadillo repeat domain (ARM), followed by two tandem sterile alpha motif domains (SAM), and a toll-interleukin receptor homology domain (TIR) that possess the NAD^+^ glycohydrolase (NADase) activity required for axon degradation. SARM1 is the founding member of a family of TIR-domain containing enzymes that catalyze the cyclization reaction of NAD^+^ to produce ADP-ribose (ADPR) and cyclic ADP-ribose (cADPR). Its activation leads to cellular NAD^+^ depletion, resulting in metabolic failure and axon fragmentation^3–6^. Loss of SARM1 in animal models protects axons in various neurodegenerative conditions including chemotherapy-induced peripheral neuropathy (CIPN), traumatic central and periphery nerve injury, and glaucoma^7–13^. Additionally, activating mutations in SARM1 have been found in patients with amyotrophic lateral sclerosis (ALS)^14–16^, indicating a general and conserved neural degenerative function of the protein.

The SARM1 ARM domain contains an allosteric pocket in which the binding of NMN disrupts the ARM-TIR interactions, leading to TIR domain self-association and the formation of functional catalytic sites capable of cleaving NAD^+^^17–19^. Conversely, NAD^+^ can bind to the same site, competing with NMN to prevent SARM1 activation^20,21^. Thus, the cellular ratio of NMN/NAD^+^, which is increased due to the diminishing of a fast-turnover NMN to NAD^+^ converting enzyme NMNAT2 (Nicotinamide Nucleotide Adenylyltransferase 2) in the proximal portion of the injured nerves, has been proposed to be the key determinant controlling SARM1 activation^22^. The NMN/NAD^+^ ratio model of SARM1 activation is supported by the fact that NMN accumulates prior to axon degeneration after injury, and SARM1 can be directly activated by adding NMN to purified SARM1^19,23^. However, the concentration of NMN required for SARM1 activation *in vitro* exceeds that of the cellular concentration^24^. Moreover, in addition to axon degeneration, SARM1 activation in cultured cells causes necrotic cell death^25^. We thus believe that the current NMN-mediated SARM1 activation model is not complete.

By observing that a group of pyridine-containing compounds, such as SIR3 (Figure 1A), activated SARM1 NADase and induced SARM1-dependent cell death and axon degeneration, we deduced a two-step activation process of SARM1 mediated by those compounds. In the first step, SARM1 catalyzes a base exchange reaction between the SARM1-activating compounds and the nicotinamide moiety of NAD^+^, resulting in the formation of ADPR conjugates. Then the Compound-ADPR conjugates induce SARM1 condensates formation as it promotes the self-proliferating assembly of parallel, intertwined superhelical structures in which numerous functional catalytic sites of the TIR domains are stably formed. This mechanism reconciles the previously proposed NMN activating model and provides a mechanistic explanation for how SARM1 activation is spatially restricted to damaged axons.

**Figure 1.**
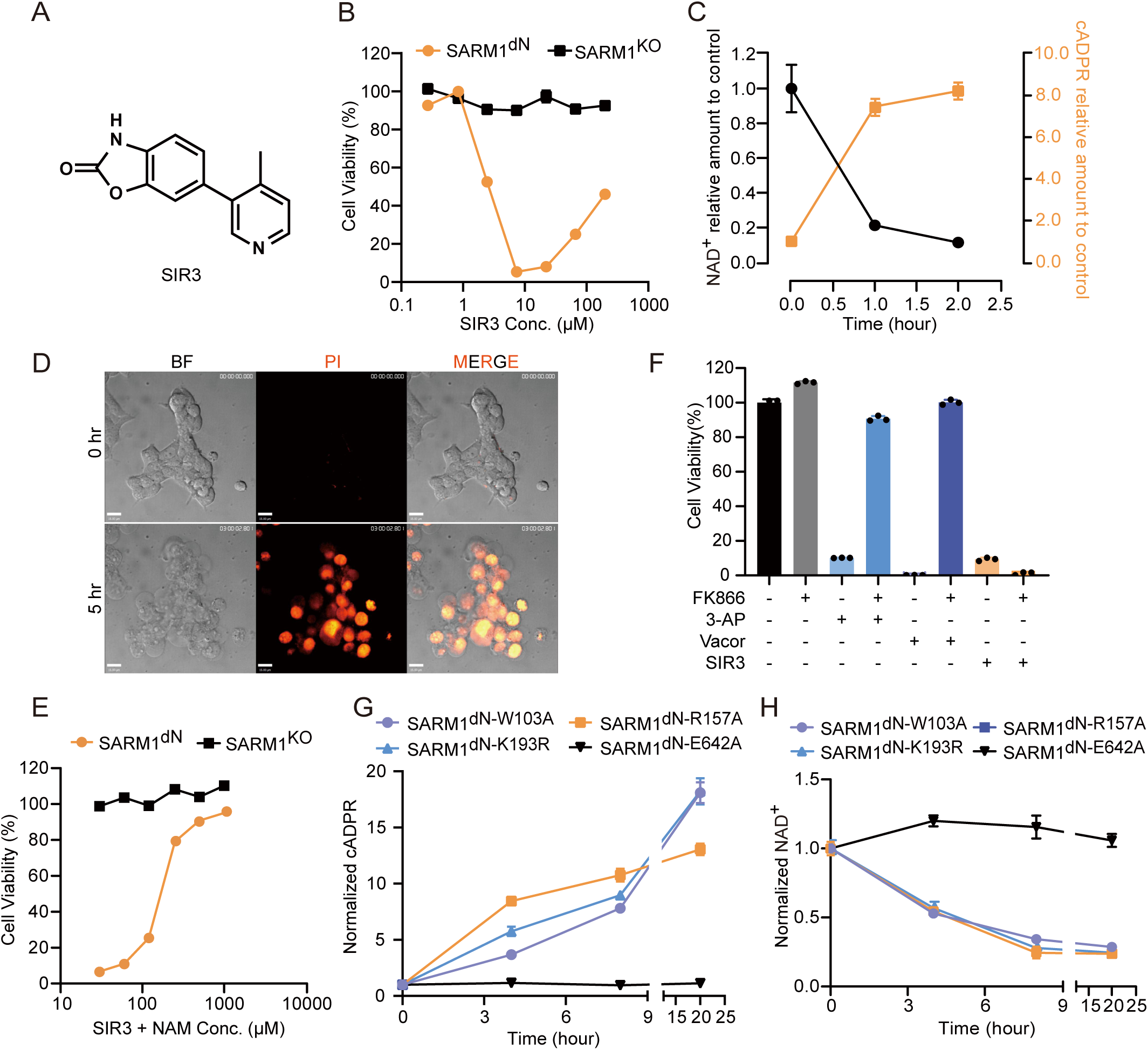
SIR3 triggers SARM1-dependent cell death (A) Chemical structure of the hit compound, SIR3. (B) SARM1^-/-^ and SARM1^dN^-overexpressing HEK-293T cells were treated with varying concentration of SIR3 as indicated. Cell viability was measured with the CellTiter-Glo assay. (C) SARM1^dN^-overexpressing HEK-293T cells were treated with SIR3 (7 μM). NAD^+^ and cADPR were quantified by LC/MS at indicated times and normalized to 0 hr after treatment. (D) SARM1^dN^-overexpressing HEK-293T cells were treated with SIR3 (7 μM) for 5 hours and cell membrane disruption was visualized with propidium iodide (PI) staining. Scale bars, 15 μm. (E) SARM1^dN^-overexpressing HEK-293T cells were treated with SIR3 (7 μM) plus different concentration of NAM as indicated. Cell viability was measured with the CellTiter-Glo assay. (F) SARM1^dN^-overexpressing HEK-293T cells were treated with FK866 (200 nM), 3-AP (100 μM), Vacor (10 μM), SIR3 (7 μM) and a combination of FK866 with 3-AP, Vacor, or SIR3 for 12 hours. Cell viability was measured with the CellTiter-Glo assay. Images are representative of three independent experiments. Data with error bars are presented as the means ± SD from three independent experiments. (G and H) HEK-293T cells overexpressing SARM1 mutant (W103A, R157A, K193R, and E642A) were treated with SIR3 (7 μM). Metabolites (G) cADPR and (H) NAD^+^ were quantified by LC/MS at indicated times after treatment. Images are representative of three independent experiments. Data with error bars are presented as the means ± SD from three independent experiments. Statistical analysis was performed using Student’s t test.

Unexpectedly, we found that this activation mechanism is shared by a class of potent SARM1 NAD^+^ hydrolase inhibitors currently under clinical development, including DSRM-3716, NB-3, NB-7, and related molecules^26–28^, raising concerns about their clinical usage.

## Results

### SIR3 induces SARM1 activation differently from NMN mimics

To investigate the cell death mechanism mediated by SARM1, a HEK-293T cell line with its endogenous SARM1 knocked out was used to stably express SARM1 (dN-SARM1) lacking the mitochondrial localization sequence (residues 1-27). The morphology and proliferation rate of those cells were similarly to that of the parental HEK-293T cells (Figure S1A). When treated with the NMN-mimicking SARM1 activator CZ-48 or NMN mimic metabolic precursors 3-AP and Vacor^29,30^, SARM1-dependent cell death occurred (Figures S1B-1D). Using this cell line for a screening assay for similar cell killing chemicals, we identified a novel compound SIR3 that promoted SARM1-dependent cell death as well (Figure 1A). Interestingly, the SARM1-dependent cell death induced by SIR3 was not in a linear dose-dependent fashion but rather a “U-shaped” curve (Figure 1B), contrasting with the linear dose-dependent effect observed with NMN mimetics (Figures S1B-1D). Notably, SIR3 reduced intracellular NAD^+^ while elevated cADPR levels, indicative of SARM1 NADase activation (Figure 1C)^31^. The morphology of cell death induced by SIR3 was necrotic in nature, similar to that caused by NMN mimics (Figures 1D and S1E). Also similar to the NMN mimics, the cell death was inhibited by the NADase enzymatic product, nicotinamide (NAM) (Figure 1E)^17^.

To investigate the mechanism of SARM1 activation induced by SIR3, we prepared cell lysates from the dN-SARM1 expressing cells and treated the lysates with different concentrations of SIR3 and measured the SARM1 activity with a styrylpyridine derivative PC6^32^. Interestingly, unlike NMN, adding SIR3 to the cell lysates was unable to activate SARM1 directly (Figure S1F). One possibility for this observation is that SIR3 needs to be metabolized into a NMN mimic similar to 3-AP and Vacor. Both compounds are metabolized to 3-AP mononucleotide (3-APMN) and Vacor mononucleotide (VMN) by nicotinamide phosphoribosyl transferase (NAMPT) before gaining ability to bind and activate SARM1. As such, the NAMPT inhibitor FK866 protects cells against 3-AP- and Vacor-mediated cell toxicity^29,30^. However, treatment of cells with FK866 did not block SIR3-induced cell death but rather promoted it (Figures 1F and S1G). It is worth noting that unlike in cultured cells, co-treatment of SIR3 and FK866 of cell lysates still failed to activate SARM1 (Figure S1H). It is thus clear that SIR3-induced SARM1 activation and ensuing cell death is different from that induced by 3-AP and Vacor.

All the previously known SARM1 activators, including NMN, CZ48, 3-APMN, and VMN, activate SARM1 by binding to the ARM domain allosteric pocket of SARM1^22^. Mutations in this domain including W103A, R157A, and K193R rendered the host cells resistant to those NMN mimics (Figures S1I-S1K). However, treatment of those cells with SIR3 still caused a significant increase in cellular cADPR levels and decrease in NAD^+^ (Figures 1G and 1H).

Overall, our data suggest that SIR3-induced activation of SARM1 was inconsistent with the NMN/NAD^+^ ratio model with NMN binding to the ARM domain of the allosteric pocket to achieve SARM1 NADase activation.

### SIR3-activated SARM1 forms condensates in cells

To dissect the molecular mechanism underlining SIR3-induced SARM1 activation, we treated HEK-293T cells stably expressing dN-SARM1 with SIR3 for indicated time and then fractionated the cell lysate into different cellular fractions through differential centrifugation before subjecting them to PC6-based NADase assay (Figure S2A). As shown in Figure 2A, compared to the vehicle (DMSO-treated) cells, the SARM1 activity was detected in the P13 fraction (13000 *g* centrifugation pellet) of the SIR3-treated cells, and the activity in the P13 fraction could not be further activated by NMN, indicating the SARM1 in the P13 fraction was already fully activated by the SIR3 treatment. Since such a fractionation procedure normally collected heavy membrane organelles like mitochondria and lysosomes in P13, it was a surprise that SIR3-activated SARM1 ended up there (Figure 2B). In contrast, all SARM1 protein stayed in the S13 fraction of the DMSO-treated cells and adding NMN to this fraction robustly promoted SARM1 NADase activity, confirming that SARM1 is in a much more soluble form before SIR3 treatment. A time-dependent translocation of SARM1 from the S13 (soluble) fraction to the P13 (insoluble) fraction in SIR3 treated cells further confirmed that SIR3 treatment caused SARM1 precipitation (Figure 2C). In contrast, such a SARM1 translocation was not observed after treatment with 3-AP (Figures S2B and S2C). Additionally, when subjecting the P13 fraction of SIR3-treated cells to non-reducing SDS-PAGE, large SARM1 molecular complexes were detected (Figure 2D). Those data suggested that unlike NMN mimics, SIR3 activated SARM1 by promoting the formation of large SARM1 protein complexes with active NADase activity. Those SARM1 protein complexes are “super-heavy” that they could be effectively pelleted using a centrifugation force of only 500 *g* (Figure S2D).

**Figure 2.**
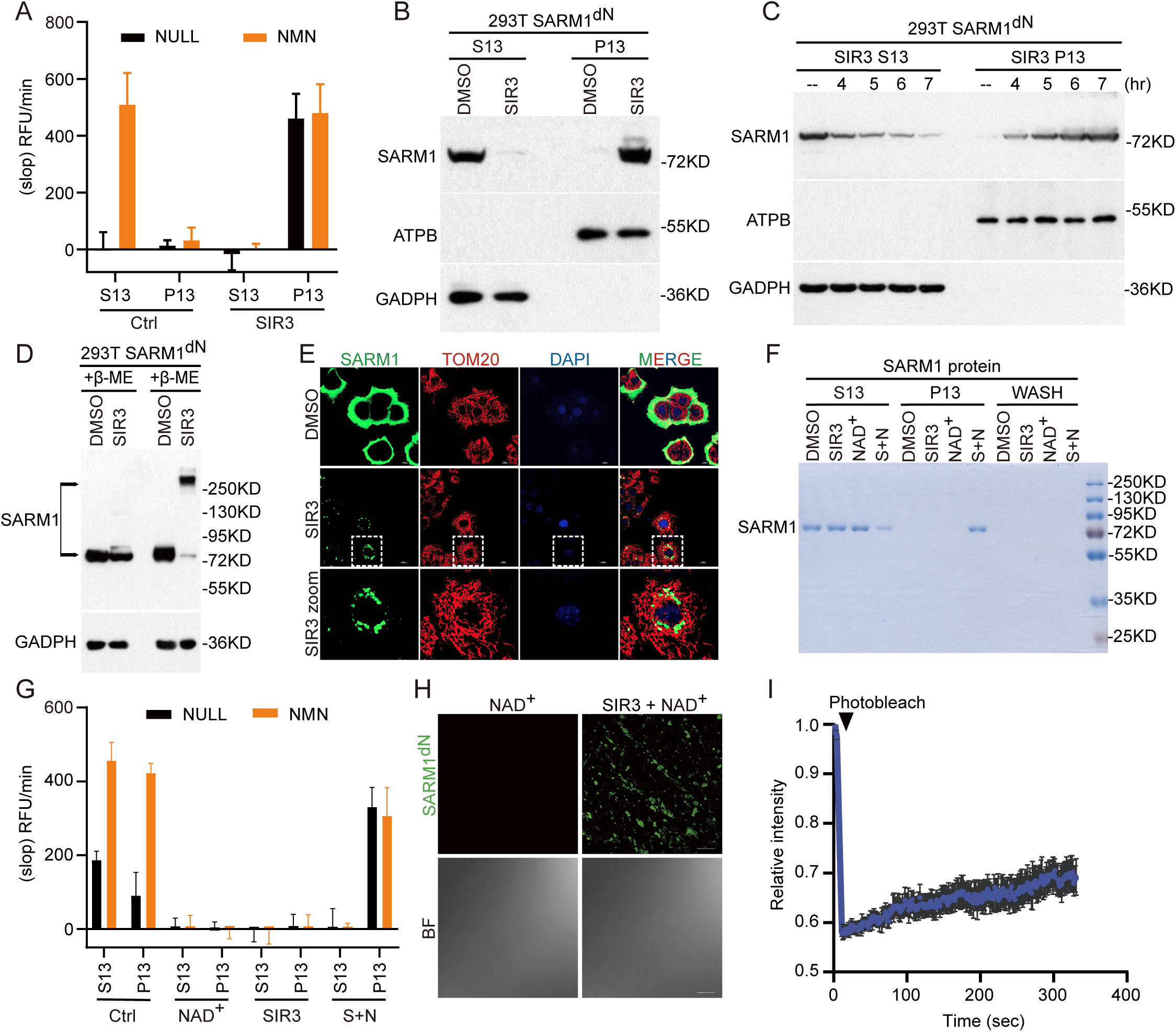
SIR3 facilitates the activation of SARM1 through a liquid-to-solid phase separation process (A and B) HEK-293T dN-SARM1 cells were treated with DMSO or SIR3, followed by lysing and centrifugation. (A) The PC6 assays were performed in both S13 (supernatant) and P13 (precipitant) in the presence or absence of 200 μM NMN. (B) The western blots were also performed in both S13 and P13 using the indicated antibodies. (C) HEK-293T dN-SARM1 cells were treated with SIR3 for the indicated time period, followed by lysing and centrifugation, the supernatant and precipitant fractions were subjected to western blot to assess SARM1 levels. (D) Cell lysates were immunoblotted under non-reducing conditions using the indicated antibodies. (E) HEK-293T dN-SARM1 cells were treated with SIR3 (7 μM) for 4 hours and, then fixed, and stained for SARM1 (anti-SARM1 antibody, Green), TOM20 (anti-TOM20 antibodies, red), and chromosome (DAPI, Blue). Scale bars as indicated. (F and G) Purified dN-SARM1 protein were treated with DMSO, NAD^+^ (100 μM), SIR3 (100 μM) and SIR3 + NAD^+^ for 1 hour in vitro followed by centrifugation. After that, both S13 and P13 were applied to (F) coomassie blue staining and (G) the PC6 assay in the absence or presence of 200 μM NMN. (H) SIR3/NAD^+^ induces liquid-to-solid transition of the GFP-SARM1 proteins. SIR3/NAD^+^ was incubated with GFP-SARM1 at 37 °C for 1 hour in vitro. Scale bars, 40 μm. (I) Dynamics analysis of GFP-SARM1 aggregates by FRAP. Images or Blots are representative of three independent experiments. Data with error bars are presented as the means ± SD from three independent experiments. Statistical analysis was performed using Student’s t test.

To visualize those SARM1 complexes in cells, we performed immunostaining of SIR3-treated cells using antibodies against SARM1 and other organelle markers. The result showed that SARM1 formed puncta structures in those cells after treated with SIR3. Notably, these cytoplasmic puncta did not co-localize with TOM20, LAMP1, DAPI, the respective cellular marker of mitochondrial, lysosome, and nuclear (Figures 2E and S2E). In contrast, those SARM1 condensates were not detected in cells treated with 3-AP (Figure S2F). The SARM1 condensates were also detected in the cytosol of SIR3-treated cells as clustered gold particles by immune-electron microscopy (Figure S2G).

Consistently, the allosteric pocket mutants of SARM1 still formed active condensates in response to SIR3 treatment. The TIR mutant, E642A, did not show such a response (Figures S2H-S2L). These results further indicate that SIR3 activates SARM1 differently than all the previously reported NMN mimics.

### SIR3-induced SARM1 activation through a NAD^+^-dependent liquid-to-solid phase transition

We further attempted to reconstitute SIR3-induced SARM1 activation *in vitro*. To our surprise, SIR3 alone could not activate and precipitate SARM1 in the cell lysate like it did in cells (Figure 3A). However, in the present of NAD^+^, SIR3 robustly activated SARM1 in the cell lysate by promoting the formation of active condensates, which could not be further activated by NMN (Figures S3B and S3C). These data indicated that the SIR3-mediated SARM1 activation was driven by a phase transition process, and NAD^+^ is essential for such reaction. The SIR3-induced SARM1 condensates collected by 13,000 *g* centrifugation effectively preserved the NAD^+^ hydrolyzing activity *in vitro.* Even after 24 hours incubation at 37, the NADase activity of SARM1 condensates remained unchanged, while the SARM1 in the S13 fraction from the vehicle-treated cells completely lost its ability to response to NMN after 5 hours, indicating that once activated by SIR3, SARM1 remained in a very stable form (Figures S3D-3F). Similarly, SIR3 precipitated and activated purified SARM1 in the present of NAD^+^ (Figures 2F and 2G). Once precipitated, the SARM1 condensates maintained their active form even after extensive washing with a buffer no longer containing SIR3 and NAD^+^, confirming that the condensates, once formed with their NADase activated, were irreversible. To visualize the condensates directly, we further purified recombinant GFP-fused SARM1 protein and activated it with NAD^+^ and SIR3 before observation under a confocal microscope. The image showed the clumps of GFP-SARM1 protein in the presence of NAD^+^ and SIR3 (Figure 2H). Consistent with this observation, FRAP assays demonstrated relatively low dynamics of GFP-SARM1, indicating that GFP-SARM1 proteins formed solid-phase condensates (Figures 2I and S3G). In addition, 1,6-hexanediol, a compound able to break reversible liquid-to-liquid phase transition, did not affect the condensation and activity of SARM1 (Figures S3H and S3I)^33–35^. All these experiments suggest that SIR3 activates SARM1 through an irreversible liquid-to-solid phase transition.

**Figure 3.**
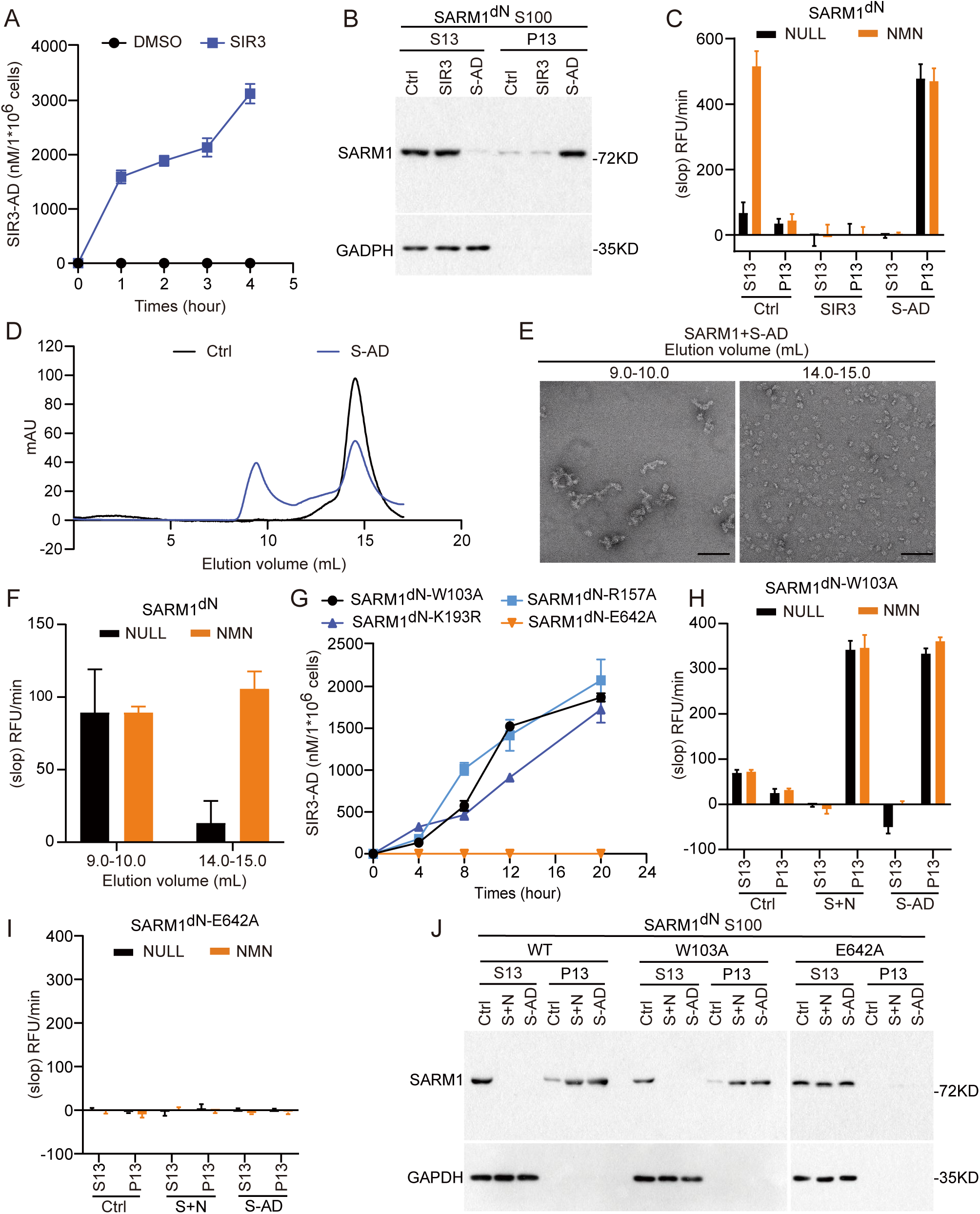
SIR3-ADPR adduct was identified as the genuine activator of SARM1. (A) Production of SIR3-ADPR in SARM1^dN^-overexpressing HEK-293T cells treated with SIR3 (7 μM). SIR3-ADPR were quantified by LC/MS at indicated times after treatment. (B and C) dN-SARM1 containing lysates were treated with DMSO, SIR3 (100 μM) and SIR3-AD (100 μM) for 1 hour in vitro, followed by centrifugation. After that, both S13 and P13 were applied to (B) western blots and (C) the PC6 assay in the absence or presence of 200 μM NMN. (D) Analytical size-exclusion chromatography analyses of dN-SARM1 and dN-SARM1+SIR3-ADPR (100 μM). Curves are representative of three independent experiments. (E) Representative EM negative stain image of SARM1 dN-SARM1 incubated with SIR3-ADPR (100 μM) from (D). Scale bars, 100 nm. (F) The PC6 assays were performed in fractions from (E) in the presence or absence of 200 μM NMN. (G) Production of SIR3-ADPR in SARM1 W103A, R157A, K193R, and E642A mutants’ cells. (H-J) SARM1 WT, W103A, and E642A mutant’s lysate were treated with DMSO, SIR3 (100 μM) + NAD^+^ (100 μM) or SIR3-ADPR (100 μM), followed by centrifugation. After that, both S13 and P13 were applied to (H and I) the PC6 assay in absence or presence of 200 μM NMN and (J) western blots using the indicated antibodies. Images or Blots are representative of three independent experiments. Data are presented as the means ± SD. n = 3 independent experiments. Statistical analysis was performed using Student’s t test.

### SARM1-catalyzed NAD^+^ base exchange is required for SARM1 activation

In addition to its NAD**^+^** hydrolysis activity, the same enzymatic site of SARM1 is able to catalyze base-exchange between the nicotinamide moiety of NAD**^+^** and molecules containing pyridyl group to form an ADPR-conjugates (Figure S4A)^36^. Since SIR3 is a pyridyl-group-containing molecule and its activation of SARM1 requires the presence of NAD^+^, we thus hypothesized that the SARM1 base-exchange product SIR3-ADPR-conjugates but not SIR3 itself was the molecule that directly induced SARM1 phase transition and activation. Indeed, significant amount of SIR3-ADPR was observed in a time-dependent fashion in cells treated with SIR3 (Figure 3A). To verify this hypothesis directly, we generated and purified SIR3-ADPR, then added it into the SARM1-expressing cell lysate. Indeed, the SARM1 in the lysate underwent phase transition and activation, a process no longer required the presence of additional NAD^+^ (Figures 3B, 3C and S4B, S4C). Meanwhile, when analyzed by a non-reducing SDS-PAGE, SARM1 protein was found to be in high molecular weight complexes after incubation with SIR3-ADPR, similar to what happened to SARM1 when SARM1-expressing cells were treated with SIR3 (Figures 2D and S4D). These experiments indicated that once SIR3-ADPR conjugate is formed, it is sufficient to activate SARM1 through phase transition.

We subsequently subjected the purified SARM1 incubated with SIR3-ADPR to analytical size-exclusion chromatography. Compared with SARM1 alone, SARM1 incubated with SIR3-ADPR eluted from the column as high molecular weight aggregates (Figure 3D). Furthermore, electron microscopy images showed that these aggregates exhibited a distinct filament morphology, unlike the octameric structure of native SARM1 (Figure 3E). Notably, unlike the SARM1 octamers that can be activated by NMN, those high molecular weight SARM1 aggregates induced by SIR3-ADPR was already in a fully activated state and cannot be further activated by NMN (Figures 3F and S4E). These findings suggest that SIR3-ADPR-conjugates directly activate SARM1 by inducing the formation of large molecular condensates.

Moreover, the generation of SIR3-ADPR conjugate was detected in cells expressing SARM1 NMN/NAD^+^ allosteric binding pocket mutants following the treatment with SIR3 (Figure 3G). Therefore, it is not surprising that SIR3-ADPR still induced SARM1 W103A condensation and activation (Figures 3H and 3J). This finding further validated that the induction of SARM1 phase transition by SIR3-ADPR is independent of the NMN/NAD^+^ allosteric binding pocket. Conversely, SIR3-ADPR did not induce the condensation of SARM1 E642A mutant (Figures 3I and 3J), indicating that the E642 site of SARM1 is not only crucial for its base-exchange and NADase activity but also essential for phase transition induced by SIR3-ADPR.

### SIR3-induced cell death of endogenous SARM1-expressing cells and axonal degeneration requires a priming step

Surprisingly, unlike HEK-293T cells ectopically expressing dN-SARM1, the parental HEK-293T cells and the dorsal root ganglia (DRG) neurons isolated from mice were not able to undergo cell death and axonal degeneration when treated with SIR3 (Figures S5A-S5C).

To study the reason behind such a difference, we first generated a HEK-293T cell line stably expressing full length SARM1 (FL-SARM1) with the endogenous SARM1 knocked out, allowing better control of SARM1 expression levels. Interestingly, even when FL-SARM1 was expressed at the similar level as dN-SARM1, the treatment of those cells with SIR3 did not induce much cell death (Figures 4A and S5D), despite a decrease of NAD^+^ and increase of cADPR were detected in the treated cells (Figures S5E and S5F). Furthermore, the generation of SIR3-ADPR in those FL-SARM1 expressing cells was also observed, indicating the occurrence of the base-exchange reaction (Figure S5G). However, the amount of SIR3-ADPR generated in FL-SARM1 expressing cells was much lower than that in cells expressing dN-SARM1 under the same treatment duration (Figure 3A). This discrepancy can be attributed to the lower basal base-exchange activity of FL-SARM1 compared to dN-SARM1 (Figures 4B, S5H and S5I). This observation suggested that in order to achieve full activation of FL-SARM1 by SIR3, a “priming” event, i.e., increased SARM1 NAD^+^ hydrolysis and base exchange activity may be required for the generation of sufficient SIR3-ADPR for the condensate formation and activation for the native FL-SARM1 whereas SARM1 without its mitochondrial targeting sequence (dN-SARM1) already has enough basal level activity to by-pass this step. To test this hypothesis directly, we co-treated those cells with sub-lethal concentrations of CZ48 or 3-AP with SIR3 and observed a substantial incidence of cell death. Again, the cell death induced by CZ48/3-AP and SIR3 displayed a U-shape concentration curve (Figures 4C and S5J-S5L).

**Figure 4.**
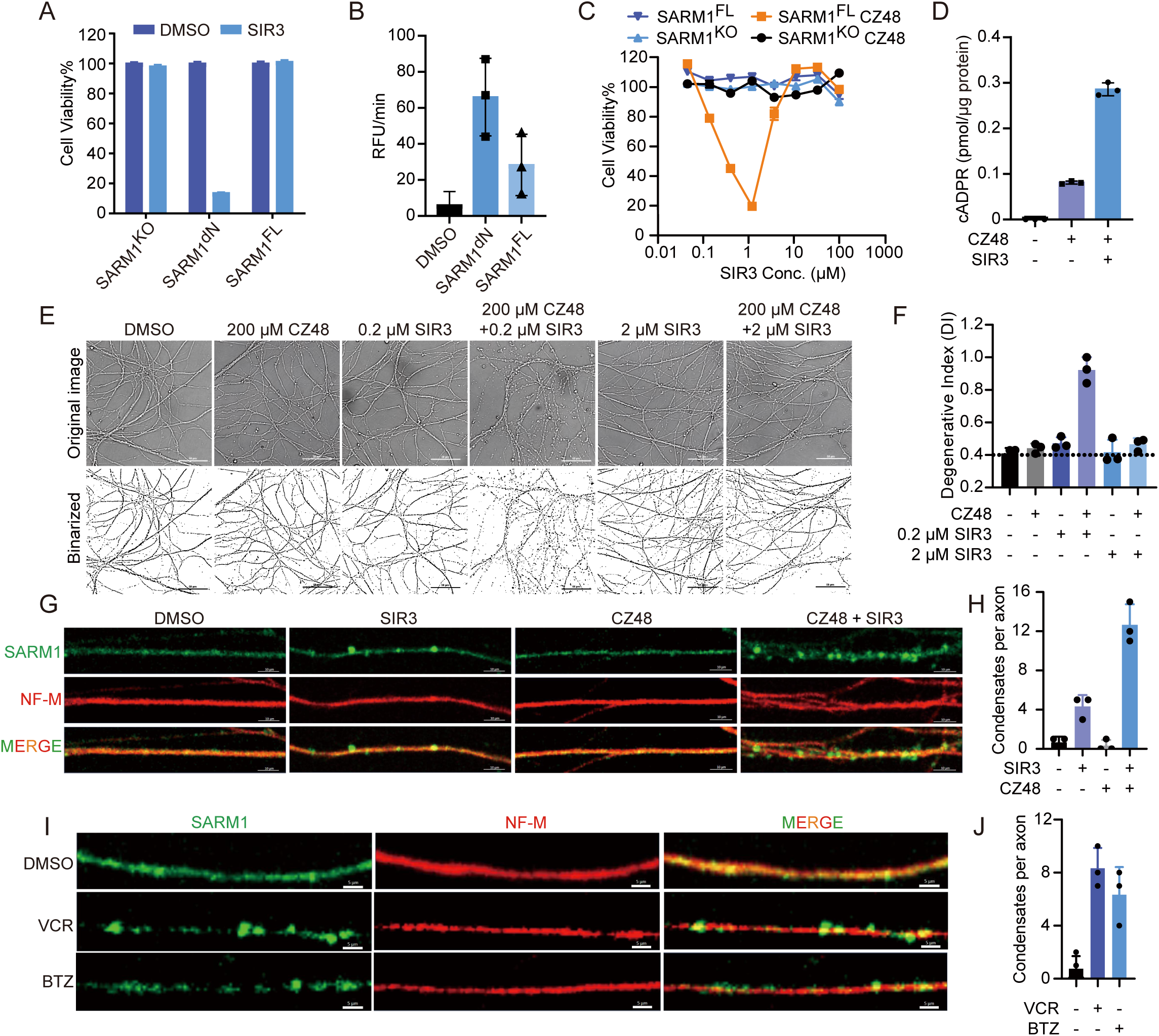
SIR3 can induce SARM1-dependent axon degeneration. (A) SARM1^-/-^, 293T dN-SARM1 and 293T FL-SARM1 were treated with DMSO, SIR3 (7 μM) for 12 hours. Cell viability was measured with the CellTiter-Glo assay. (B) SARM1 activity in lysates with the same concentrations of 293T dN-SARM1 and 293T FL-SARM1 was evaluated by PC6 assay. (C) 293T SARM1^−/−^ and FL-SARM1 cells were treated with CZ48 (10 μM) or a combination of CZ48 with different doses of SIR3 as indicated. Cell viability was measured with the CellTiter-Glo assay. (D) cADPR level from wild-type embryonic DRG neurons treated with CZ48 (200 μM) or a combination of CZ48 with SIR3 (0.2 μM), measured by LC-MS/MS. (E and F) Representative images of embryonic DRG neuron axons after treatment with different compounds, as indicated, for 72 hours. Binarized images (ImageJ) are displayed to enhance the visualization of axonal integrity during the analysis of the degeneration index. (F) Quantification of DRG axonal degeneration corresponding to (E). Scale bars, 50 μm. (G) Immunofluorescent staining with antibodies against SARM1(green) or neurofilament (red) of embryonic DRG neuron axons from WT mice in which the DRG were treated with DMSO, SIR3 (0.2 μM), CZ48 (200 μM) or a combination of CZ48 with SIR3. Scale bars, 10 μm. (H) Number of SARM1 condensates per axon area from (G). (I) Immunofluorescent staining with antibodies against SARM1(green) or neurofilament (red) of embryonic DRG neuron axons from WT mice in which the DRG were treated with DMSO, Vincristine or Bortezomib (BTZ). Scale bars, 10 μm. (J) Number of SARM1 condensates per axon area from (I). Images are representative of three independent experiments. Data are presented as the means ± SD. n = 3 independent experiments. Statistical analysis was performed using Student’s t test.

Interestingly, 3-AP alone was also unable to induce cell death in the parental HEK-293T cells even at 10 mM concentration compared to 1/100 that amount required for SIR3 priming (Figures S5M and S5N). Similarly, in DRG neurons, treatment with CZ48 alone, even at a concentration as high as 400 μM, which exceeds the cellular NMN level by two orders of magnitude, did not cause axonal degeneration (Figures 4D, S5O and S5P)^22^. However, when co-treated with SIR3, CZ48 and 3-AP both induced a robust axonal degeneration in DRG neurons and cell death in HEK-293T cells. Notably, the axonal degeneration and cell death also followed a U-shaped concentration curve of SIR3 (Figures 4D-4F and S5N). DRG neurons isolated from the SARM1 knockout mice were totally resistant to such treatment, indicating that axon degeneration in DRG neurons induced by CZ48 and SIR3 was SARM1-dependent (Figures S5Q AND S5R). The SARM1 in the DRG neurons treated with CZ48 and SIR3 showed obvious condensates, whereas treatment with SIR3 alone resulted in fewer condensates, and CZ48 treatment showed no observable condensates when immune-stained with an anti-SARM1 antibody (Figures 4G and 4H). SIR3, therefore, is able to induce cell death of endogenous SARM1-expressing cells like the parental HEK-293T cells and axonal degeneration of DRGs, but doing so needs a priming event carried out by NMN mimics.

Moreover, chemotherapy-induced peripheral neuropathy (CIPN) is a common complication during cancer treatment. Previous studies have indicated that axonal degeneration induced by two commonly used chemotherapeutic agents, vincristine (VCR) and bortezomib (BTZ), is attributed to SARM1 activation^37,38^. Indeed, we observed the formation of SARM1 condensates in isolated mouse DRG neurons treated by these chemotherapy drugs, further emphasizing the significance of SARM1 condensate formation in its activation (Figures 4I and 4J).

### The SARM1 condensates are parallel, intertwined superhelixes

As shown in Figure 2H, the precipitated SARM1 condensates exhibited clump aggregates under microscope. However, these images of the precipitated SARM1 condensates did not give much structural information about how those condensates were formed and how they remain functional as NADase. To address this question, we subjected what still remained in the soluble fraction after purified SARM1 protein was treated with SIR3-ADPR, a fraction should contain the intermediates of those higher-order of condensates, to cryo-EM analysis. Previous size-exclusion chromatography confirmed that SARM1 in those fractions were large complexes with active NADases activity (Figure 3F). The 2D classification of those intermediates showed that SARM1 protein bound to SIR3-ADPR possessed the curved double-stranded TIR domains with the octameric SAM or SAM-ARM rings attached to the convex surface (Figures S6A and 6B). From the dataset of SARM1 bound to SIR3-ADPR, we obtained a cryo-EM structure of the double-stranded octameric TIR domain with well-defined density for SIR3-ADPR at 3.85 Å resolution (Figures 5A-5C, S6B, S6C and S7A). The two strands are arranged in an anti-parallel direction, with each strand aligned in a head to tail fashion (Figure 5B). SIR3-ADPR occupies the catalytic site and function as a molecular glue to facilitate the assembly of each strand (Figures 5D, 5E and S7B). In addition, SIR3-ADPR strengthens the interactions between the two strands by forming a hydrogen bond with E686 on the EE loop of the opposite strand via the benzoxazolinone group that is attached to the pyridine (Figure 5E). This group also makes a hydrogen bond with N679 on the EE loop. In contrast, these hydrogen bond interactions are absent for 1AD or NAD^+^.

**Figure 5.**
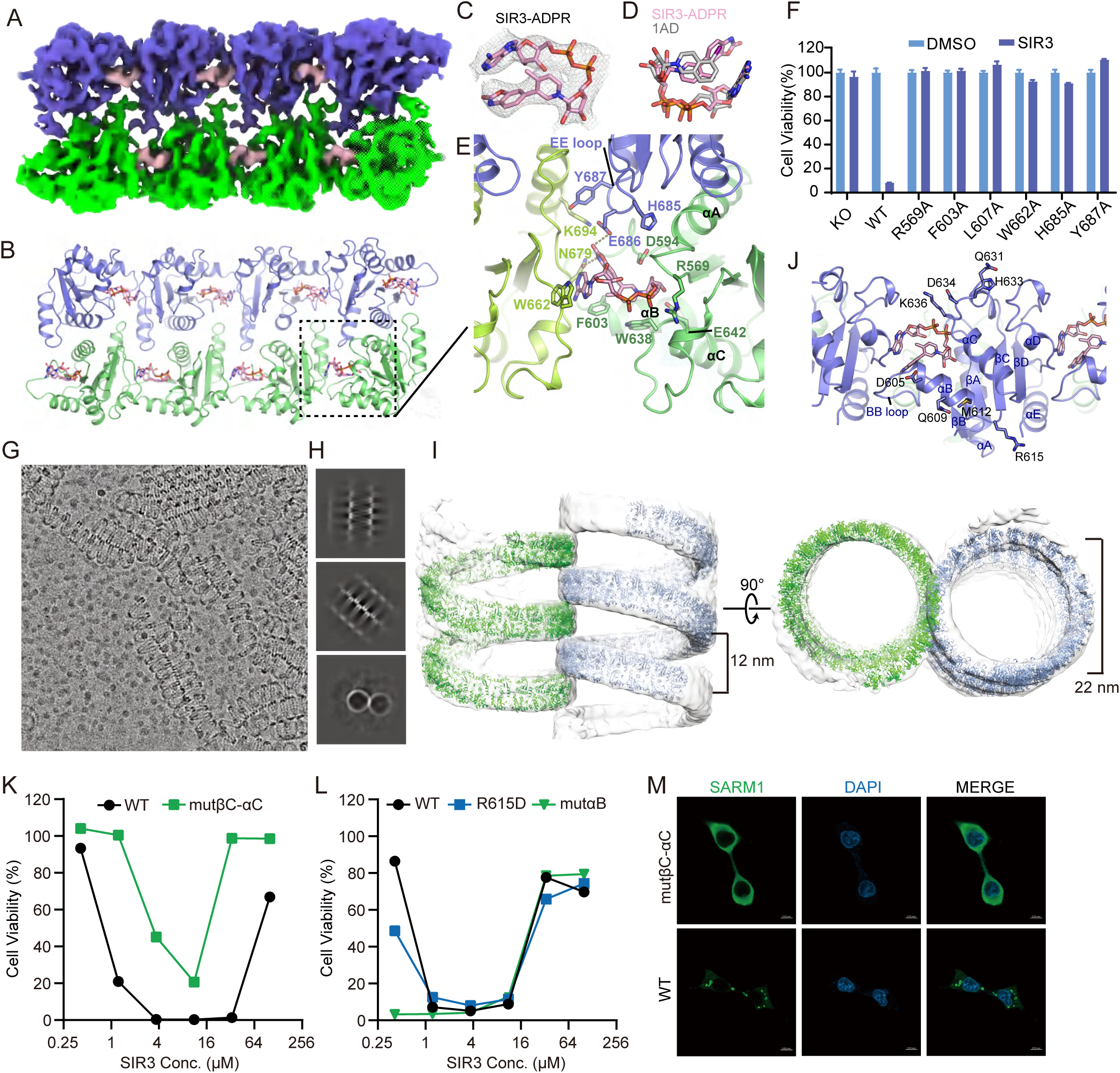
SARM1 condensates involve the assembly of intertwined superhelical structures. (A) Cryo-EM map of the SARM1 TIR domain bound to SIR3-ADPR reconstructed from the dataset of SARM1 FL bound to SIR3-ADPR. SAM and ARM domains are not visible due to their high flexibility relative to the TIR domain. (B) Cryo-EM structure of the SAMR1 TIR domain bound to SIR3-ADPR. (C) EM density map for SIR3-ADPR. (D) Superposition of the structures of SIR3-ADPR and 1AD in the TIR domain. (E) Detailed interactions between SIR3-ADPR and the TIR domain. (F) SARM1^−/−^, SARM1^−/−^ 293T cells stably expressing SARM1 orthostatic-site mutations were treated with DMSO or SIR3 for 12 hours. Cell viability was measured with the CellTiter-Glo assay. (G and H) Representative cryo-EM micrograph (G) and 2D class averages (H) of the TIR domain of SARM1 in complex with SIR3-ADPR. (I) Cryo-EM map of the TIR domain bound to SIR3-ADPR reveals two parallel intertwined superhelical structures. Structure of the octameric TIR domain bound to SIR3-ADPR (b) are docked in the map. (J) Residues that are potentially involved in interactions between two neighboring superhelical structures are shown as sticks. The orientation of the model is orthogonal to that shown in (B). (K and L) Effects of mutations in the TIR domain on SARM1-mediated cell death. SARM1^−/−^ 293T cells stably expressing SARM1 WT or mutants were treated with increasing concentrations of SIR3. Cell viability was measured with the CellTiter-Glo assay. Data are presented as the means ± SD from three independent experiments. (M) Mutations of residues possibly involved in the formation of intertwined superhelical structures prevent condensates formation in cells. Images are representative of three independent experiments. Data are presented as the means ± SD. n = 3 independent experiments. Statistical analysis was performed using Student’s t test.

To further confirm the importance of SARM1/SIR3-ADPR interaction in the condensate formation and activation of SARM1, we systematically mutated those ADPR conjugates binding site and measured the effect on SARM1 activation. The alanine mutations of R569, that interacts with the diphosphates of SIR3-ADPR; F603 and L607 that interact with the pyridine moiety; and W662 that interacts with the adenine moiety, all abolished the ability of SIR3-ADPR to induce the condensate formation and activation of SARM1 as well as cell death (Figures 5E, 5F and S7B-S7F). Furthermore, the residues H685 and Y687 in the EE loop, which are involved in two-stranded TIR domain assembly (Figures 5E and S7B), are crucial for the activation of SARM1 by ADPR conjugates (Figures S7G and S7H). These findings validated the essentiality of SIR3-ADPR binding in inducing condensate formation and activation of SARM1.

To further elucidate the higher-order oligomerization structure of the TIR domain induced by SIR3-ADPR, we purified the TIR domain of SARM1 and subject it to cryo-EM analysis. As expected, addition of SIR3-ADPR into the purified TIR domain resulted in the detection of a superhelical filamentous structure with variable helical pitch, indicating the conformational flexibility of the oligomerization interface of the TIR domain (Figure 5G). Interestingly, the superhelical structures are intertwined with each other and appeared in parallel arrays (Figures 5G and 5H). Single particle cryo-EM analysis yielded the reconstructions of the TIR domains with two parallel, intertwined superhelical structures at about 15 Å overall resolution (Figure S6C). Despite the low resolution due to conformational heterogeneity, the cryo-EM structure of the double-stranded TIR domain bound to SIR3-ADPR could be docked into the map (Figure 5I). The overall structure is characterized by a left-handed superhelical solenoid with 22 nm diameter and a 12 nm helical pitch. 48 TIR monomers are present in each turn of the superhelical structure. Therefore, we reasoned that the formation of intertwined superhelical structure of the TIR domain could facilitate condensate formation and contribute to SARM1 activation. Notably, the intertwined superhelical structure was also observed in the FL-SARM1 bound to SIR3-ADPR (Figure S6A). Due to the resolution limitations, the inter-helical contact cannot be precisely located but is limited to the solvent-exposed residues on αB, αB βC loop, βC αC loop or βD αD loop (Figure 5J). While mutations of D605, Q609 and M612 (mutαB) on αB or R615 on the αB βC loop showed minimal effect on SIR3-mediated cell death, mutations of Q631, H633, D634 and K636 (mutβC αC) on the βC αC loop partially but not completely inhibited SIR3-induced cell death and prevented condensation process in cells (Figures 5K and 5M). This is likely because the formation of individual superhelical solenoids is sufficient to activate SARM1, while interactions between multiple superhelical solenoids can further promote SARM1 protein phase separation, enhancing the reaction rate. Taken together, the intertwined superhelical structures of SAMR1 bound to SIR3-ADPR are critical in determining phase separation of SARM1 that leads to the formation of numerous functional catalytic sites for NAD^+^ hydrolysis.

### Inhibitors of SARM1 TIR domain activate its NADase through phase transition

The fact that loss-of-enzymatic-function mutants of SARM1 confer neuroprotection has prompted great interest in developing SARM1 enzyme inhibitors for the treatment of neurodegenerative disorders^39,40^. A class of potent and selective pyridine-derived small molecule inhibitors of SARM1, targeting its TIR domain, have been recently reported with some of them advanced to phase I clinical trials^26–28^. Interestingly, all of them could be the substrates for the SARM1 catalyzed base-exchange reaction with NAD^+^. We therefore hypothesized that they also have the ability to activate SARM1 by inducing its phase transition. Indeed, molecules such as DSRM-3716, NB3, and NB7 all induced SARM1-dependent cell death at certain concentrations (Figures 6A, S8A and S8B). Notably, the induction of cell death by those compounds was also in a U-shaped concentration curve with optimal concentrations being at 4 μM, 4 μM, and 10 μM, respectively. When added to the SARM1-expressing cell lysate, all three compounds caused NAD^+^-dependent condensation and activation of SARM1 (Figures 6B, 6C and S8C-S8F). It is worth noting that continuous presence of high concentrations of those inhibitors did suppress the NADase activity of activated SARM1, effective masked their SARM1 activating capability.

**Figure 6.**
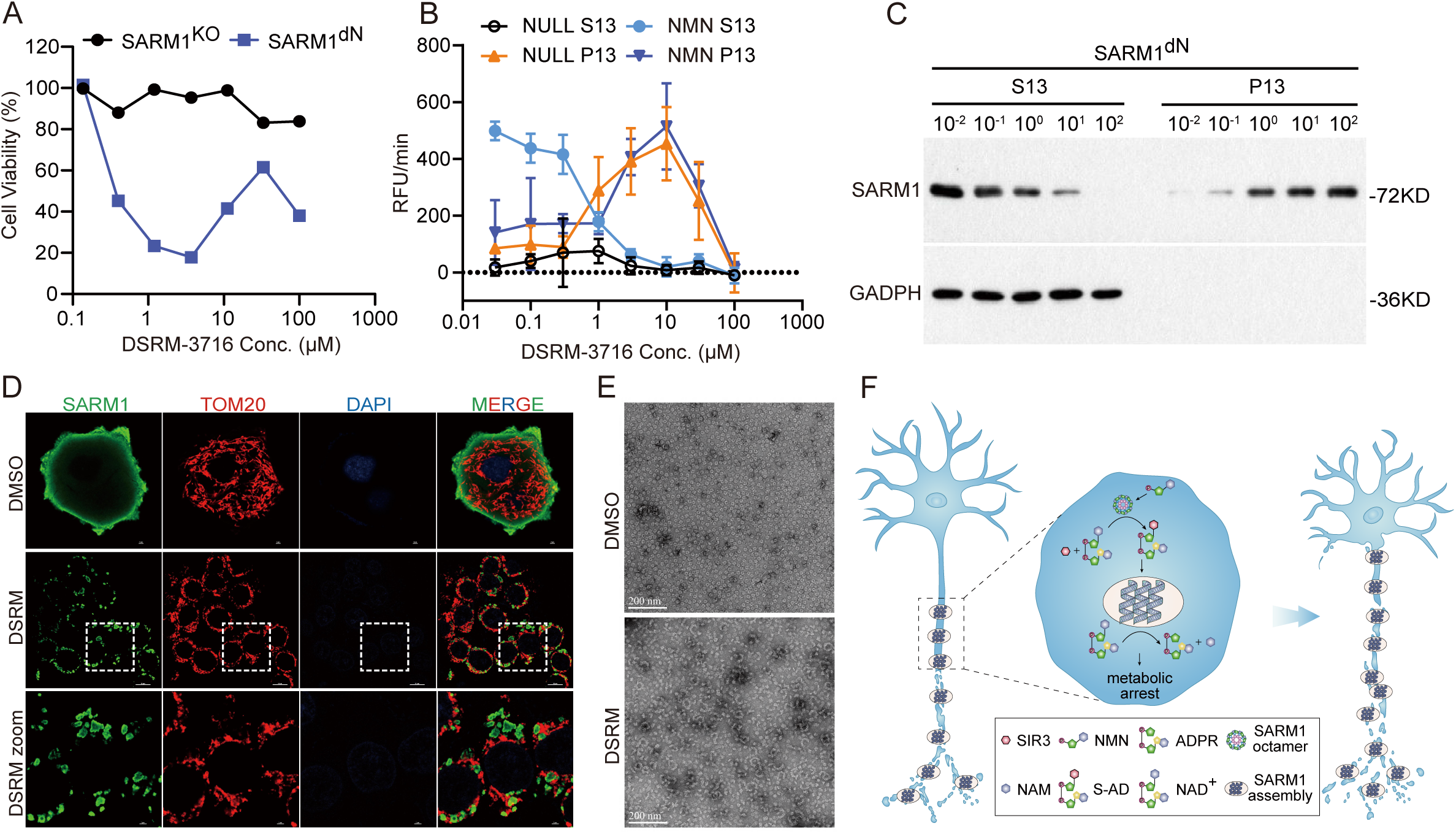
SARM1 TIR domain inhibitors have the ability to activate its NADase activity through a liquid-to-solid phase transition. (A) SARM1^dN^-overexpressing HEK-293T cells were treated with varying concentration of DSRM-3716 as indicated. Cell viability was measured with the CellTiter-Glo assay. (B and C) dN-SARM1-containing lysates were incubated with NAD^+^ (500 μM) and different doses of DSRM-3716 as indicated, followed by centrifugation. After that, (B) The PC6 assays were performed in all S13 and P13 in the presence or absence of 200 μM NMN. (C) S13 and P13 were also applied to western blots to assess SARM1 levels. Blots are representative of three independent experiments. (D) 293T dN-SARM1 cell were treated with DSRM-3716 (5 μM) for 5 hours and, then fixed, and stained for SARM1 (anti-SARM1 antibody, Green), TOM20 (anti-TOM20 antibodies, red), and chromosome (DAPI, Blue). Scale bars as indicated. (E) Representative EM negative stain image of purified dN-SARM1 incubated with DMSO or DSRM-ADPR (100 μM). Images are representative of three independent experiments. Scale bars, 200 nm. (F) Diagram of SARM1 two-step activation mechanism. Images or Blots are representative of three independent experiments. Data are presented as the means ± SD. n = 3 independent experiments. Statistical analysis was performed using Student’s t test.

We also analyzed the condensates induced by DSRM-3716-ADPR by immunofluorescent staining in cells and negative stain-EM/cryo-EM analysis in vitro. The immunostaining revealed many intracellular puncta (Figure 6D) and the negative stain-EM and cryo-EM images showed the similar structure of activated human SARM1 filaments induced by SIR3-ADPR (Figures 6E and S8G).

## Discussion

Our study has unveiled a previously unknown mechanism of SARM1 activation. During this process, pyridine-containing compounds like SIR3 are exchanged for the NAM moiety in NAD^+^ to form conjugates with the ADPR moiety of NAD^+^ by the inherent SARM1 base-exchange activity. Those ADPR conjugates, previously thought to be innocuous by-products of SARM1 NADase, have emerged as SARM1 activators. They facilitate the continuous polymerization of SARM1 proteins via their TIR domain, resulting in their transition to an active dimeric configuration and the subsequent formation of super-helical filaments with the TIR domains. The formation of those superhelical filaments is rapid and irreversible with the growth of those filaments would eventually exceed the solubility limit of the liquid phase and transit to solid phase that manifested in condensation *in vitro* and puncta formation in axons and cells. The condensation and activation of SARM1 by the pyridine-containing compounds leads to cellular NAD^+^ depletion and ensuing cell death as well axon degeneration in DRG neurons (Figure 6F). Moreover, the self-proliferating nature of superhelical SARM1 should effectively deplete the soluble SARM1 in the environment thus restricts SARM1 mediated axon degeneration locally.

In previous studies, the mechanism underlying SARM1 activation has been primarily attributed to the regulation of the NMN/NAD^+^ ratio^22^. However, it has been challenged by the fact that the concentration of NMN required for SARM1 activation *in vitro* surpasses physiological levels. Moreover, the substantial difference between NMN and NAD^+^ levels in cell, spanning over two orders of magnitude, raises concerns regarding the precise modulation of SARM1 activity through ratio manipulation^21,24^. Additionally, both our investigations and prior studies have demonstrated that administration of the NMN analogue CZ48, even at levels far exceeding cellular NMN content, fails to elicit noticeable robust axon degeneration in DRG neurons^22^. We therefore propose a two-step activation process of SARM1 uncovered by current study: the first step only requires NMN to slightly activate SARM1 activity. This activation step is not sufficient to cause cell death or axonal degeneration per se. In the second step, the NAD^+^ hydrolysis and base-exchange activity of SARM1 generates pyridine-ADPR conjugates, such as SIR3-ADPR, leading to complete and irreversible activation of SARM1 NADase through phase transition. Since the fully activated SARM1 is in a solid state, its NAD^+^ hydrolysis activity is restricted locally, a property in line with axon degeneration.

Previously, it has been reported that the SARM1-TIR domain undergoes a liquid-to-solid phase transition and activate its NADase activity in the presence of sodium citrate precipitant^41^. This study focused solely on the SARM-TIR domain and was recently challenged by other studies^27^. We also found that sodium citrate only led to a minor increase in NADase activity when dN-SARM1 was used (Figure S3J)^42^, suggesting that the phase transition to induced full activation of SARM1 is regulated by specific molecules and cannot be solely achieved by altering the environmental conditions.

Recently, it has been demonstrated that bacterial and plant TIR-domain-containing proteins exhibit NAD^+^ hydrolase activity^43,44^. Intriguing research on bacterial TIR-STING or TIR-SAVED, integral components of the cyclic oligonucleotide-based antiphage signaling system (CBASS), has provided insight into the essential role of nucleotide-regulated large molecular complexes or protein filament assembly in the activation process of these TIR enzymes^45,46^. Additionally, recent study on plant antiviral proteins containing TIR domains reveals that these proteins can induce the formation of condensates, presenting both liquid and solid phases, through their endogenous substrates NAD^+^/ATP. This phase transition process activates their NADase activity^47^. Those observation implies a highly conserved mechanism whereby small molecules to facilitate the activation of TIR enzymes NAD^+^ hydrolase through super-helical filament/condensate formation.

Finally, given the significant involvement of SARM1 in various acute and chronic neurodegenerative diseases, including CIPN, ALS, multiple sclerosis (MS), traumatic brain injury (TBI), Parkinson’s disease (PD), and Huntington’s disease (HD), coupled with the negligible impact of SARM1 knockout on normal growth and development in mice, the inhibition of SARM1 activity has emerged as an attractive therapeutic strategy for these neurodegenerative conditions^7,10,14,48–52^. However, our findings reveal an unexpected aspect: these TIR-targeting inhibitors also possess notable SARM1 activation capabilities. Moreover, although constant presence of high concentrations of inhibitors appear to fully suppress the activity of SARM1, the activation of SARM1 induced by the ADPR conjugates is irreversible. Therefore, it raises concerns regarding the potential adverse consequences arising from excessive SARM1 activation, which contradicts the intended therapeutic outcome. A recent preprint study by Genentech also observed this inhibitor-induced SARM1 activation phenomenon, although they did not elucidate its mechanism^53^. Based on the aforementioned experimental results, we rationalized that any SARM1 inhibitor generated through similar mechanisms may not completely avoid the activation side effects. This is attributable to the fact that these inhibitor-ADPR conjugates are analogs of NAD^+^, and their binding sites nearly overlap with NAD^+^ but exhibit significantly higher affinity. Consequently, these inhibitors inevitably promote conformational change and phase transition of SARM1, ultimately leading to the formation of stable and nonreversible activated SARM1 filament, whose activity will cause undesired effect, especially in neurodegenerative disease patients whose SARM1 might already be partially activated.

The current study also suggests the existence of endogenous pyridine-containing molecules that participate in the activation of SARM1 in damaged axons. Identifying those molecules could be important to understand how a variety of neuronal damaging signals activate SARM1 to cause axonal degeneration.

## Material and Methods

### Antibodies and other reagents

The antibodies used in this study included: Rabbit polyclonal anti-SARM1 antibody (abcam, ab226930), monoclonal anti-SARM1 antibody produced in mouse (made by ourselves), anti-Flag M2 antibody (Sigma-Aldrich F1804), anti-DYKDDDDK (Invitrogen MA1-91878), LAMP1 antibody (Santa Cruze sc-20011), anti-NfM (Proteintech 25805-1-AP), anti-Tom20 (Santa Cruze sc-17764), FONT-β-Actin-HRP (MBL, D291-7), anti-DDDDK-HRP (MBL, D291-7), Goat anti-Mouse IgG (H+L) Cross-Adsorbed Secondary Antibody, Alexa Fluor™ 488(Invitrogen A-11001), Goat anti-Rabbit IgG (H+L) Cross-Adsorbed Secondary Antibody, Cyanine3 (Invitrogen A10520) Chemical reagents used in this study included: DSRM-3716 (MCE, HY-W021879), CZ48 (MCE, HY-129522), 3-AP (Sigma-Aldrich, A21207), Vacor (Greyhound Chromatography), NMN (Selleck, S5259), NAD+ (Selleck, S2518), FK866 (Selleck, S2799), NAM (Sigma-Aldrich, 72340), PC6 (kind gift of SINOREX), DAPI (Invitrogen, D1306), propidium iodide (MCE, HY-D0815), Protease Inhibitor Cocktails (Merck, 04693132001)

### Cell Culture and Stable Cell Lines

HEK 293T, HEK 293T *SARM1^-/-^* and Hela cell lines were all grown in DMEM with 10% fetal bovine serum (Gibco) at 37°C under 5% CO2. All cell lines were initially purchased from the American Type Culture Collection (ATCC) and cultured according to the ATCC’s instructions. HEK-293T *SARM1^-/-^*cells were infected with virus encoding Flag tag *SARM1* (dN-SARM1 (residues 28-724) WT and mutant) and GFP-positive live cells were sorted to establish the 293T/dN-SARM1 (WT and mutant) cell lines. HEK-293T *SARM1^-/-^*cells were infected with virus encoding Flag tag *SARM1* (full length WT and mutant) and GFP-positive live cells were sorted to establish the 293T/FL-SARM1 (WT and mutant) cell lines. Hela cells were infected with virus encoding flag tag *SARM1* (dN-SARM1 (residues 28-724)) and GFP-positive live cells were sorted to establish the Hela/dN-SARM1 cell lines.

### Primary DRG cultures

C57BL/6J mice were obtained from Vital River Laboratory Animal Technology Co. and SARM1 knockout mice were generated by the animal facility of the National Institute of Biological Sciences, Beijing. All animal housed in a specific pathogen-free animal facility. Mice were housed on a 12 hr light/dark cycle with ad libitum access to food and water. Primary mouse dorsal root ganglia (DRG) was obtained as described before (Sasaki et al., 2016). Briefly, DRGs were dissected from embryonic day 13.5 mouse embryos and incubated with 0.05% Trypsin solution containing 0.02% EDTA (Gibco) at 37℃ for 30 min, and then neutralized by Fetal bovine serum. Then, DRGs was triturated by gentle pipetting and filtered through 40 µm cell strainer (Falcon). After 2 min gentle centrifugation, trypsin solution was removed and replaced for DRG growth medium (Neurobasal medium; ThermoFisher) containing 2% B27 (ThermoFisher), 50 ng/ml 2.5S NGF (Sigma), 10 μM uridine (Sigma), 10 μM 5-fluoro-20-deoxyuridine (Sigma), 50 U/ml penicillin, and streptomycin (Gibco). DRGs were resuspended in DRG growth medium at 500 DRGs/ml. The cell density of these suspensions was adjusted to 1×10^7^ cells/mL. Cell suspensions were seeded at 100,000 cells/well as a 10 μL spotted culture in the center of 24 well plates coated with poly-D-Lysine (0.1mg/mL; Sigma) and laminin (3 μg/mL; ThermoFisher), and placed in a humidified CO_2_ incubator for 15 min. After allowing for cells to adhere, culture media was added gently to a final well volume of 1 mL/well. Culture media was changed every 2 days and DRG were allowed to extend for 7 days before treatment.

## METHOD DETAILS

### DNA constructs

psPAX2, pMD2.G construct were obtained from Addgene (#12260, #12259), pX458 and pWPI construct were kept in our lab. Full-length cDNAs for human SARM1 was a gift from SIRONAX (uniprot Q6SZW1). Using this full-length cDNAs, SARM1 lacking the mitochondrial localization sequence (residues 28-724, dN-SARM1) were subcloned into pGEM-T vector and pWPI vector (GFP tagged) by using KOD polymerase (TOYOBO) to generate dN-SARM1 construct. Using Quickchange Site-Directed Mutagenesis Kit, we generate several dN-SARM1 mutants, include W103A, R157A, K193R, E642A, F603A, L607A, R569A, W662A, N679A, H685A and Y687A. In addition we also made several FL-SARM1 mutant constructs, include the full-length W103A, R157A, K193R and E642A. all the mutations were inserted into pWPI vector (GFP tagged) to generate the pWPI-dN-SARM1 (WT and mutants) and pWPI-FL-SARM1 (WT and mutants) Flag construct. 2 sgRNA targeting SARM1was subcloned to pX458 vector (PX458-GFP-SARM1#1/2).

### CRISPR/Cas9 knockout cells

To construct SARM1 knockout HEK-293T cell line, 5 μg of each pX458-GFP-SARM1 plasmid was transfected into 1*10^7^ 293T cells using the transfection reagent Lipofectamine 3000 (Thermo Fisher) following the manufacturer’s instructions. 2 days after the transfection, GFP-positive live cells were sorted into single clones by using a BD FACSFusion cell sorter. The single clones were cultured in 96-well plates for another 2 or 3 weeks or a longer time dependent upon the cell growth rate. HEK 293T *SARM1^-/-^* cell was determined by DNA sequencing and western blotting.

### Cell transfection, virus packaging and protein expression

Complementary DNA transfection was carried out using Lipofectamine3000 (Thermo Fisher) by following the manufacturer’s instructions. To prepare the virus, HEK293T cells in the 10-cm dish were transfected with 20 μg of pWPI-dN-SARM1 (WT and mutants) or pWPI-FL-SARM1 (WT and mutants) construct DNA together with 11.25 μg of psPAX2 and 3.75 μg of pMD2.G. Forty-eight hours after transfection, the lentivirus was collected, filtered through a 0.44 μm filter, and used to infect HEK 293T *SARM1^-/-^* cells. Forty-eight hours after infection, the virus-containing medium was removed and GFP-positive live cells were sorted into single clones by using a BD FACSFusion cell sorter. The protein expression was confirmed by western blotting.

### Purification of SARM1

HEK293T Flag-tagged dN-SARM1 cells were cultured to 80% confluence in 10 15-cm culture dishes. Cells were collected by scraping. The collected cell pellet was washed with cold PBS three times to remove any residual culture medium and resuspended in lysis buffer [PBS, 10% glycerol, protease inhibitor cocktail], followed by two cycles of freezing and thawing using liquid nitrogen. The cell lysate was centrifuged at 13,000 g for 30 minutes and the cleared supernatant was incubated with anti-Flag M2 agarose (60 μl) overnight at 4°C. The Flag agarose was washed with lysis buffer three times and eluted by Flag peptide (0.1 mg/ml) for 6 hours. Proteins were further purified by a Superdex 6 increase column (Cytiva) run in akta system. Peak fractions were pooled, concentrated and the purified SARM1 protein was subjected to in vitro fraction and PC6-based SARM1 NADase assays.

### Purification of SARM1-TIR

hSARM1TIR (His-Sumo-ULP1-SARM1-560-700/pET28a) were produced in E. coli (BL21C43 (DE3)) cells. Step1. Expression: hSARM1TIR-pET28a colonies were picked and grown overnight, 37℃, 200 rpm. 10 mL LB culture were inoculated to 1L LB, 37℃, 250rpm until reached to OD600=0.6-0.8, cooled to appropriate temperature(18℃), inducer (IPTG, 0.3mM) was added and grown overnight (14∼16h).

Then, cells were collect at 4℃, 4000rpm centrifuge for 10min. Step2. Purification, 1L cell was resuspended with 35ml lysis buffer (25 mM Tris, pH 7.4, 150 mM NaCl, 20 mM Imidazole), with 100U/ml DNAse I, 1000U/ml Lysozyme, 10mM MgCl2, 6mM CaCl2, 1mM PMSF and send to sonication (amplitudes 35 %, sonicate 3s, intervals 7s, total 5 min). After sonication, cell was centrifuged at 4℃, 13500rpm, for 45min, to get the supernatant and pellet. Load the supernatant collected onto the Pre-equilibrate Ni-NTA column. Wash the Ni-NTA column with 20 CVs wash buffer. ULP1 Protease was applied to the column (0.15mg ULP1 per 1L culture) and incubated overnight at 4℃, collect the flow-through sample. The target protein was concentrated and further purified by a Superdex 75 increase column (Cytiva) run in SEC buffer (25 mM Tris, pH 7.4, 150 mM NaCl, 1mM TCEP). Peak fractions were pooled, concentrated to a final concentration of approximately 3.5 mg/mL, flash-frozen as 10 μL aliquots in liquid nitrogen, and stored at −80℃.

### Preparation of SARM1 Lysate

HEK 293T dN-SARM1 (WT and mutants) or FL-SARM1 (WT and mutants) cells were cultured to confluence of 80% in 15-cm culture dishes, washed twice with cold PBS and harvested by scraping. After centrifugation at 500 x *g* for 5 minutes, the cell pellet was resuspended in 2 volumes of hypotonic buffer A [20 mM Tris-HCl pH7.5, 10 mM KCL, protease inhibitor cocktail] and put on ice. Cells were homogenized by passing the lysate through a 22 gauge needle 20 times. This cell lysate was saved as total lysate and applied to PC6-based SARM1 NADase assays. The cell total lysate was further centrifuged at 500 *g* for 10 minutes. The supernatant was centrifuged at 13,000 *g* for 10 minutes. The resulting supernatant was centrifuged again at 100,000 *g* for 60 minutes and the supernatant was saved as S100 lysate for later use in the in vitro SARM1 Lysate fraction assay.

### Cell fraction assay

HEK 293T dN-SARM1 (WT and mutants) cells were cultured to confluence of 70% in 10-cm culture dishes, treated by different compounds for 4-7 hours, after treatment, the cell were lysed and centrifuged at 13,000 *g* for 10 minutes, the supernatant (S13) and precipitant (P13) fractions were collected and applied to western blotting and PC6-based SARM1 NADase assays.

### SARM1 Lysate or purified SARM1 protein fraction assay

dN-SARM1 S100 lysate with a final concentration of 2 mg/ml or purified SARM1 protein with a final concentration of 0.2 mg/ml was incubated with the respective compound in the present of 10 mM MgCl_2_ at 37 ℃ for indicated time, DSRM-3716 for 1 hour or NB3/NB7 for 3 hours. After incubation, the S100 lysate or purified SARM1 protein were further centrifuged at 13,000 *g* for 10 minutes, the supernatant (S13) and precipitant (P13) fractions were collected and applied to western blotting/coomassie blue staining and PC6-based SARM1 NADase assays. The supernatant S13 and P13 fractions of purified SARM1 were also applied to electron microscopy negative stain assay.

### PC6-based SARM1 NADase assays

The reaction buffer includes 10 μM PC6, 250 μM NAD with/out 200 μM NMN in Dulbecco’s PBS buffer in a final assay volume of 60 μl. Kinetic fluorescence reading (Ex/Em: 390 nm/510 nm) was immediately started after mixing the lysate containing SARM1 or purified SARM1 protein with the reaction buffer in a 96-well microplate, reading in the Infinite M200 PRO microplate reader (Tecan). The slopes of initial fluorescence kinetics (RFU/min) were calculated for the NADase activities of SARM1.

### FRAP assay

FRAP of SARM1-TIR condensates in vitro was conducted with samples in slices using Olympus FV4000 confocal microscope using a ×40 objective. Puncta were bleached with a 488 nm laser pulse. The recovery time was recorded for the indicated time as mentioned. Image analysis was made by Cellsene software. Data were plotted by using Prism8.

### Immunofluorescent staining

Hela flag-tagged dN-SARM1 Cells were seeded on poly-D-lysine - coated cell culture dish (NEST 801001) and treated with DSRM-3716. Wild-type and SARM^-/-^ DRG neurons were isolated and seeded as described above, and treated with CZ-48 and/or DSRM-3716 at DIV 7. The cells were immunostained as the following procedure. The cells were fixed in 4% paraformaldehyde (BIORIGIN) for 30 minutes followed by gentle PBS-T rinse, and permeabilized with 0.1% Triton X-100 in PBS for 30 min. After blocking with skim milk (5% in PBST) for 30 minutes, the cells were incubated overnight in primary antibody at 4°C. The following primary antibodies were used: anti-NfM (Proteintech 25805-1-AP; 1: 300), anti-flag (Invitrogen MA1-91878; 1:200), anti-Tom20 (Santa Cruze sc-17764; 1:200), and anti-SARM1 (home-made; 1:200). Samples were gently washed 3 times with PBST. The fluorescence was developed by incubating with Alexa Fluor 488 or Cyanine3-conjugated secondary antibody (Invitrogen A-11001/A10520; 1:300), and images were acquired with a confocal microscope (Nikon A1) and a super-resolution microscope (Nikon N-SIM) with a 20× object (for DRG neurons) and a 100× oil object (for Hela cells). NIS-Elements AR analysis (Nikon) software were used for the image analysis.

### Quantification of axon degeneration

DRG neurons were cultured in 24 well plates with cell bodies sequestered to allow imaging of axons by automated microscopy. Bright-field images per well of live axons were acquired at indicated time points using a confocal microscope (Nikon A1) with a 20× object (for DRG neurons). Axon degeneration index, which is quantified based on axon morphology using an ImageJ-based script, has been extensively described elsewhere (Sasaki et al., 2009a).

### Metabolite extraction and measurement

Intracellular metabolites were extracted by addition of 1mL iced (vol:vol) 70% acetonitrile in 30% ddH_2_O. After brief vortexing and incubation on ice for 5 min, samples were centrifuged at 4 °C for 15,000 [rpm 10 min to pellet precipitated proteins. The clear aqueous phase was transferred into a microfuge tube and lyophilized under vacuum. Lyophilized samples were stored at −20 °C until measurement. Metabolites (1AD, NAD^+^, cADPR) were measured using LC-MS/MS as described previously (<u>Shi</u> et al.<u>, 2022, Molecular Cell 82, 1643–1659</u>)

### Electron microscopy negative stain assay

The 200 mesh formvar-carbon coated EM grids (Beijing XXBR Technology Co.,ltd; T10044) were Glow-discharged for 1 min by plasma cleaner to increase their hydrophilicity. Then, 5 μL dN-SARM1 (0.2 mg/ml) Proteins were deposited on formvar-carbon coated EM grids for 30 sec. After side blotting, the grid was immediately stained with 2% uranyl acetate (UA) and then blotted again from the side. Staining was repeated twice with a 30 sec incubation with 2% UA in the final staining step. EM images were collected on a TECNAI spirit G2 (FEI; Eindhoven, Netherlands) transmission electron microscope at 120 kV.

### Immunoelectron microscopy (IEM)

Hela dN-SARM1 cells grown on sapphire discs were cryo-immobilized by HPF (HPF COMPACT 01, Engineering office M. Wohlwend; Sennwald, Switzerland) and freeze-substituted in anhydrous acetate containing 0.1% UA. Then the cells were infiltrated and embedded in LR White resins according to the instructions. 90 nm thick slices were mounted on nickel grids for immunolabeling. Grids were incubated in solutions by floating section side down on a drop of solution and transferred sequentially from drop to drop to complete the immunolabeling. To cover a grid, 5–50 μL drops of solution are placed on a piece of Parafilm in a Petri dish kept in a moist chamber to avoid drying of the solution. Sections were incubated in blocking buffer (2% (w/v) bovine serum albumin in PBS buffer) for 10 min and then incubated overnight at 4℃ in a moist chamber with primary antibody (Monoclonal anti-Flag M2 antibody produced in mouse, F1804, Merck) diluted in blocking buffer. Following 5×1 min washes in PBS buffer they were incubated in species-specific 10 nm gold conjugate (polyclonal goat anti-mouse antibody, G7652, Sigma) diluted 1:100 in blocking buffer. Following washing buffer and distilled water washes, the grids were pecessed by gold enhance kit (GoldEnhance-EM plus 2114, Nanoprobes, Inc.) to enlarge the colloidal gold particles for 3 min. After rinsing with distilled water, the sections were contrasted with uranyl acetate and lead citrate. Images were aquired on a TECNAI spirit G2 (FEI; Eindhoven, Netherlands) transmission electron microscope at 120 kV.

### Cryo-EM specimen preparation and data collection

To prepare the specimen of SARM1 (TIR)-SIR3-ADPR complex, 1mM SIR3-ADPR was added to the SARM1 (TIR) protein sample and incubated for 0.5 to 1 hour. Glow-charged 300-mesh holey carbon grids (Quantifoil R1.2/1.3 Au) were loaded into a Vitrobot Mark IV chamber (Thermo Fisher Scientific) set at 8°C in temperature and 100% in humidity. 3.0 μl of sample was applied to the grid, blotted for 5.0 s and plunged into liquid ethane for rapid freezing. All the grids were stored in boxes in liquid nitrogen prior to Cryo-EM data collection. Cryo-EM movies were captured using EPU on a Titan Krios G4 microscope (Thermal Fisher Scientific) equipped with a Falcon 4 direct electron detector operating at 300 kV and ×96,000 magnification with a pixel size of 0.808 Å.

### Cryo-EM data processing

A total of 3,567 movie stacks were subjected to MotionCor2 for motion-correction, followed by patch-based CTF estimation in CryoSPARC. A total of 383,397 particles were picked with blob picker, extracted with a box size of 600 pixels and subjected to 2D classification, which yielded two major classes: ring-like structure and superhelical structures. Due to the large size of the superhelical structures, we picked particles using Template Picker with the superhelical structure as a template and extracted particles with a larger box size of 1,200 pixels. The particles were binned 4 times to a box size of 300 pixels. Two rounds of 2D classification were performed to select particles with the superhelical structure. Finally, 69,935 particles were subjected to ab-initio reconstruction. The particles showing superhelical structures were re-extracted with a box size of 1,200 pixels and binned 2 times to 600 pixels, and subjected to heterogeneous refinement, resulting in a final map of intertwined superhelical structure with a global resolution of 15.5 Å.

### Measuring cell survival and cell death

Cell survival was measured with Cell Titer-Glo (Promega, G7570) luminescent viability assay according to the manufacturer’s instructions. Membrane leakage of necroptotic cells was measured by Cyto Tox-Glo (Promega, G9292) luminescent assay according to the manufacturer’s instruction. Briefly, cells were plated in 96-well culture plate at 5000 cells per well 24 hours before the assay. After compounds treatment, plates were allowed to reach room temperature. A total of 30 μl of either CellTiter-Glo or CytoTox-Glo reaction solution was added into each well and mixed with cell culture medium by pipetting. The reaction was carried out on a horizontal shaker at room temperature for 15 minutes. Luminescent signals were measured in a plate reader (PerkinElmer, EnSpire). Necrotic cell death was also measured in live cells by applying propidium iodide in culture medium and imaging by spinning disk confocal microscope.

### Statistical analysis

Statistical analysis was performed with Prism8 software (GraphPad). Enzyme reaction rate, intracellular metabolite content, cell death or cell survival rate were shown as means ± SD. Differences between means were assessed by Student’s t test (two-tailed, unpaired), ∗∗∗p < 0.001.

## Supplemental figure legends

**Figure S1.**
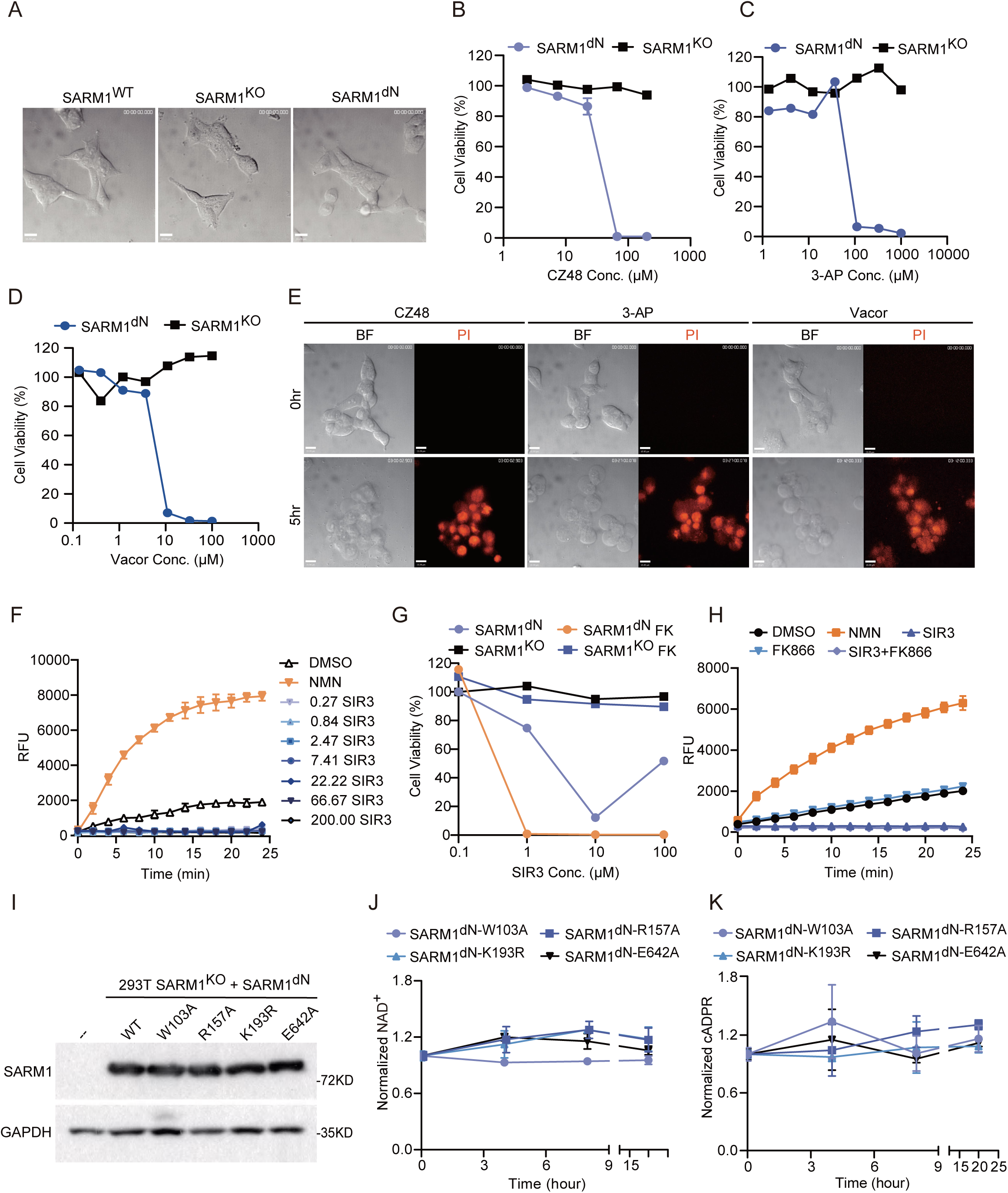
SIR3-induced SARM1-dependent cell death was different from NMN mimics, related to Figure 1. (A) The morphology of HEK-293T WT, SARM1^−/−^, and SARM1^dN^-overexpressing cells. Scale bars, 15 μm. (B-D) SARM1^dN^-overexpressing HEK-293T cells were subjected to treatment with varying concentrations of (B) CZ48, (C) 3-AP, and (D) Vacor, as indicated. Cell viability was measured with the CellTiter-Glo assay. (E) SARM1^dN^-overexpressing HEK-293T cells were treated with CZ48 (100 μM), 3-AP (100 μM), Vacor (10 μM) for 8 hours and cell membrane disruption was visualized with propidium iodide (PI) staining. Scale bars, 15 μm. (F) 293T dN-SARM1 cell lysates (2 mg/mL) were incubated with NMN and different doses of SIR3 as indicated, NADase activities were measured by PC6 assay. (G) SARM1^dN^-overexpressing HEK-293T cells were treated with indicated concentrations of SIR3 (low, medium, and high) as well as FK866 in combination with these varying concentrations of SIR3. Cell viability was measured with the CellTiter-Glo assay. (H) 293T dN-SARM1 cell lysates were incubated with DMSO, NMN (200μM), FK866 (100μM), SIR3 (100μM), or FK866 + SIR3, NADase activities were measured by PC6 assay. (I) Western blot of the expression of dN-SARM1 and mutants (W103A, R157A, K193R, and E642A) in HEK-293T SARM1^-/-^ cells. (J and K) SARM1 mutant cells were treated with CZ48 (100 μM). Metabolites (J) NAD^+^ and (K) cADPR were quantified by LC/MS at indicated times after treatment. Images or Blots are representative of three independent experiments. Data are presented as the means ± SD. n = 3 independent experiments. Statistical analysis was performed using Student’s t test.

**Figure S2.**
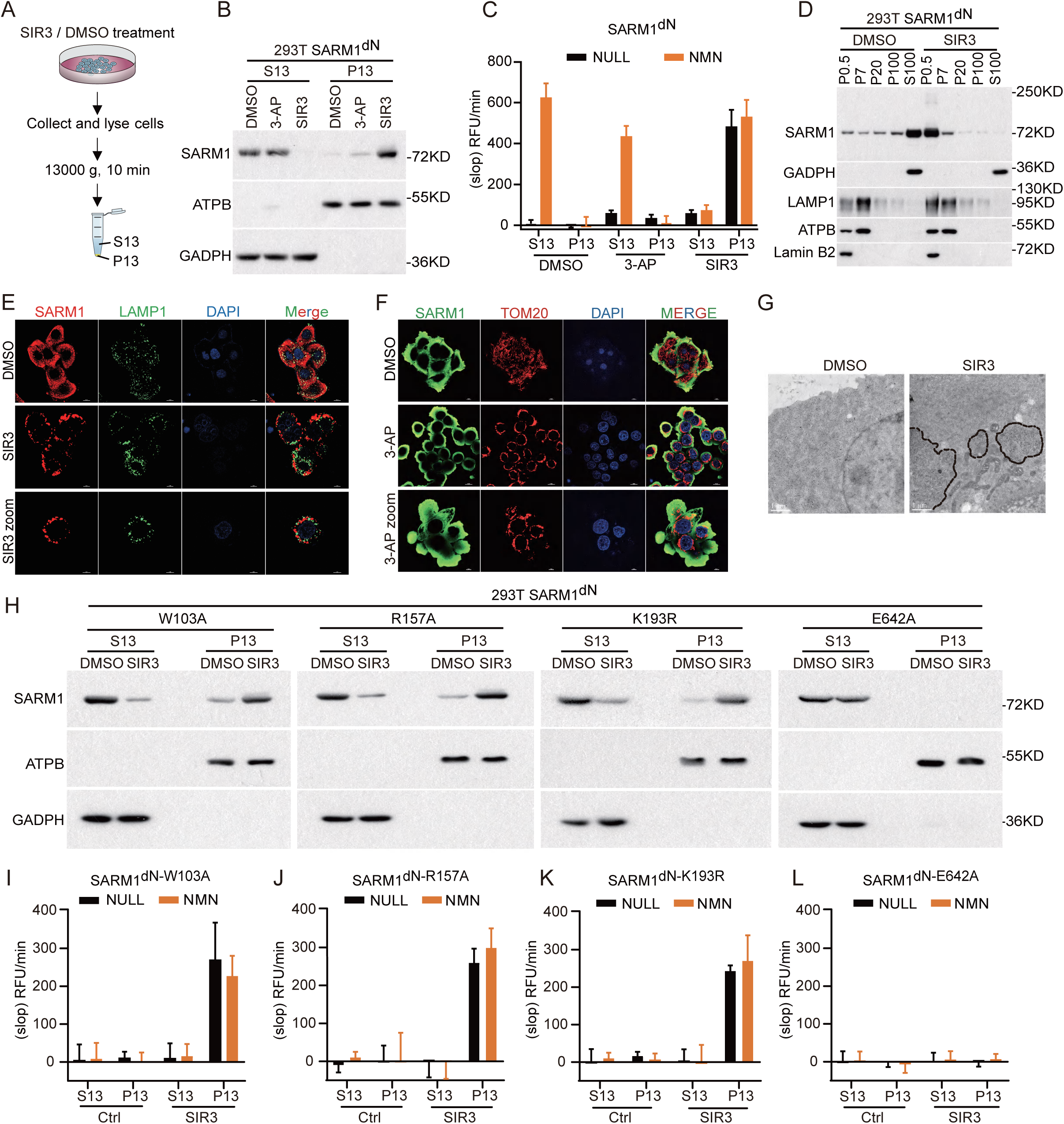
SIR3-induced SARM1 phase separation, related to Figure 2. (A) A schematic diagram of the integration of cellular assays and in vitro PC6 NADase assays. (B and C) HEK-293T dN-SARM1 cell were treated with DMSO, 3-AP (100 μM) or SIR3 (7 μM), followed by lysing and centrifugation. (B) The western blots were performed in both S13 and P13 using the indicated antibodies. (C) The PC6 assays were also performed in both S13 and P13 in the presence or absence of 200 μM NMN. (D) HEK-293T dN-SARM1 cell were treated with DMSO or SIR3 (7 μM) for 7 hours. After treatment, cell fractionation was performed by differential centrifugation, followed by western blot analysis using the indicated antibodies. (E) HEK-293T dN-SARM1 cells were treated with SIR3 (7 μM) for 4 hours, then fixed, and stained for SARM1 (anti-SARM1 antibody, Green), LAMP1 (anti-LAMP1 antibodies, red), and chromosome (DAPI, Blue). Scale bars as indicated. (F) HEK-293T dN-SARM1 cells were treated with 3-AP (100 μM) for 6 hours, then fixed, and stained for SARM1 (anti-SARM1 antibody, Green), TOM20 (anti-TOM20 antibodies, red), and chromosome (DAPI, Blue). Scale bars as indicated. (G) Cells were immunolabeled with M2-FLAG antibodies, followed by observation of the subcellular localization of SARM1 using electron microscopy. Scale bars, 1 μm. (H) SARM1 mutant cells (W103A, R157A, K193R, and E642A) were treated with SIR3 (7 μM), followed by lysing and centrifugation, the supernatant and precipitant fractions were subjected to western blot to assess SARM1 levels. (I-L) The PC6 assays were also performed in both S13 and P13, in the presence or absence of 200 μM NMN, in SARM1 (I) W103A, (J) R157A, (K) K193R and (L) E642A mutants. Images or Blots are representative of three independent experiments. Data are presented as the means ± SD from three independent experiments. Statistical analysis was performed using Student’s t test.

**Figure S3.**
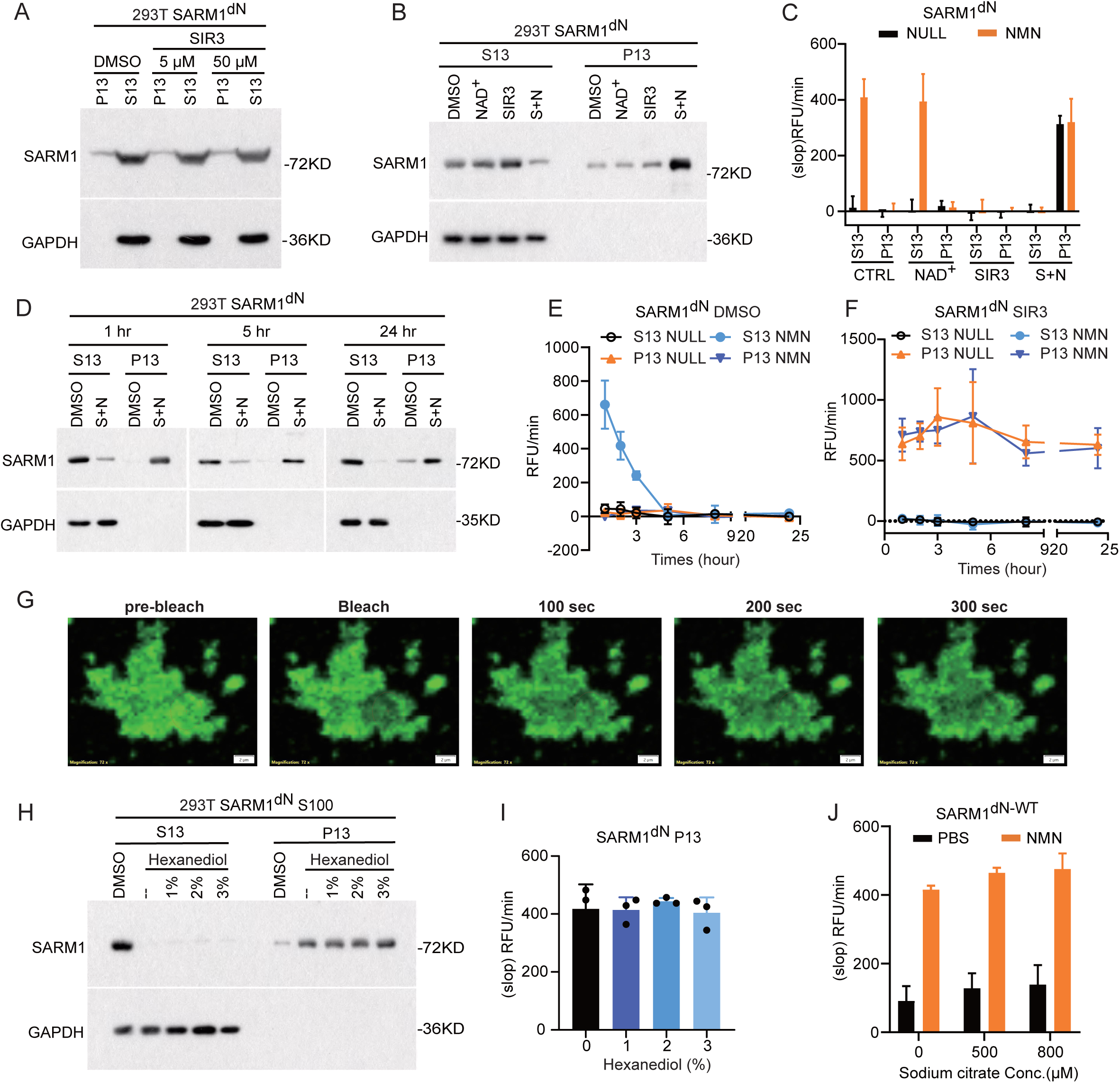
SIR3 triggers the formation of active SARM1 condensates, related to Figure 2. (A) dN-SARM1 containing lysates were treated with DMSO or difference concentration of SIR3 as indicated for 1 hour in vitro, followed by centrifugation. After that, both S13 and P13 fractions were applied to western blots. (B and C) dN-SARM1 containing lysates were treated with DMSO, NAD^+^ (100 μM), SIR3 (100 μM) and SIR3 + NAD^+^ for 1 hour in vitro, followed by centrifugation. After that, both S13 and P13 were applied to (B) western blots using the indicated antibodies and (C) the PC6 assay in the absence or presence of 200 μM NMN. (D-F) dN-SARM1 containing lysates were treated with DMSO or SIR3 + NAD^+^ for the indicated time period, followed by centrifugation, the S13 and P13 fractions were subjected to (D) western blot and (E-F) the PC6 assay in the absence or presence of 200 μM NMN. (G) Image of GFP-SARM1 aggregates by FRAP at indicated time points. Scale bar = 2 μm. (H and I) dN-SARM1 containing lysates were treated with DMSO, SIR3 + NAD^+^ and a combination of SIR3 + NAD^+^ with difference concentration of 1,6-hexanediol for 1 hour in vitro, followed by centrifugation. After that, both S13 and P13 were applied to (H) western blots using the indicated antibodies and (I) the PC6 assay in the absence or presence of 200 μM NMN. (J) Purified dN-SARM1 protein were incubated with different concentration of sodium citrate as indicated for 30 minutes in vitro. NADase activities were subsequently measured using the PC6 assay in the absence or presence of 200 μM NMN. Images or Blots are representative of three independent experiments. Data are presented as the means ± SD from three independent experiments. Statistical analysis was performed using Student’s t test.

**Figure S4.**
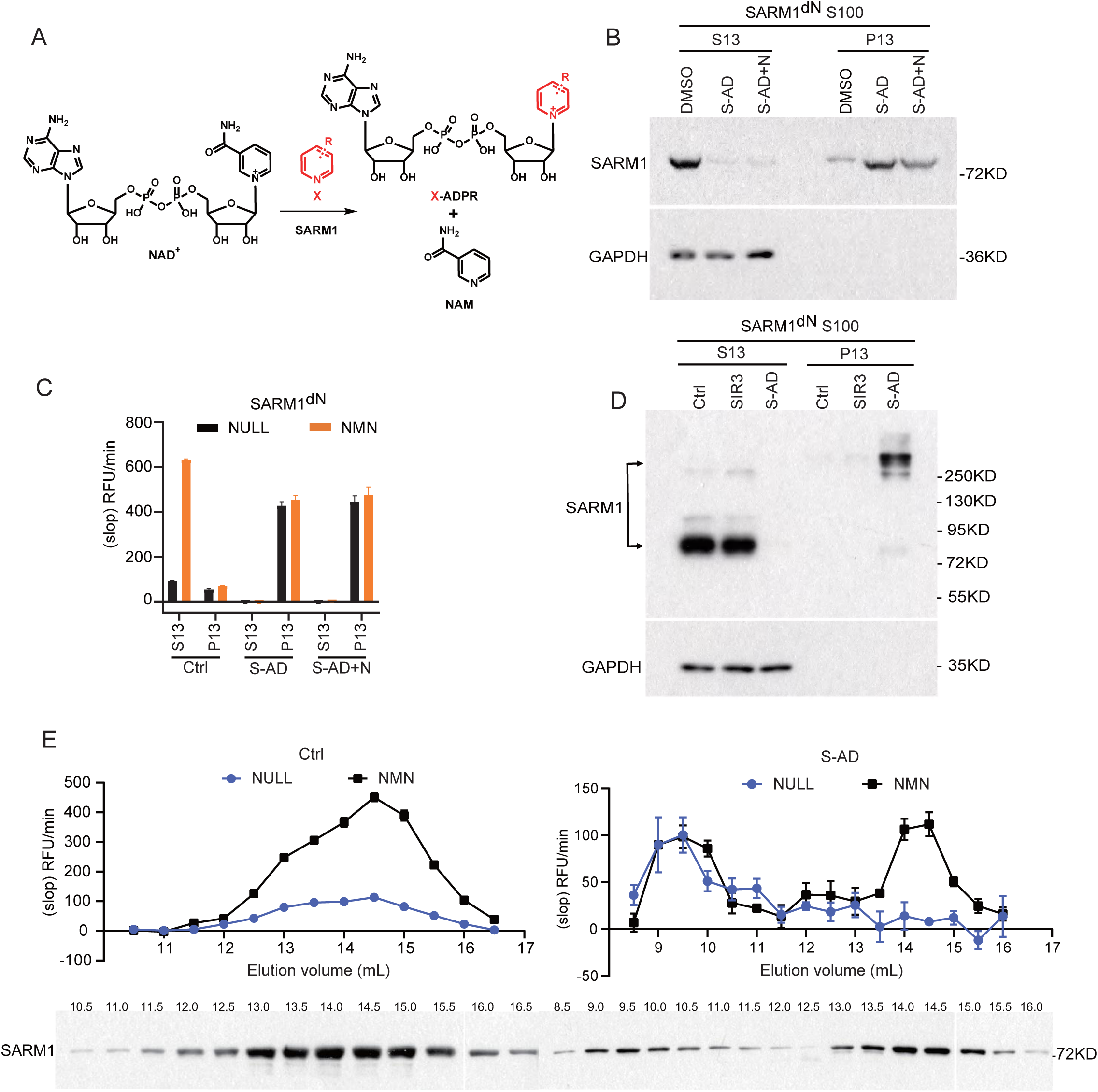
SARM1 base exchange product is the *bona fide* activator, related to Figure 3. (A) SARM1 base change reaction replacing NAM with pyridine-containing molecule. (B and C) dN-SARM1 containing lysates were treated with DMSO, SIR3-ADPR (100 μM) and SIR3-ADPR + NAD^+^ (100 μM) for 1 hour in vitro, followed by centrifugation. After that, both S13 and P13 were applied to (B) western blots and (C) the PC6 assay in absence or presence of 200 μM NMN. (D) dN-SARM1 containing lysates were treated with DMSO, SIR3 (100 μM) and SIR3-ADPR (100 μM) for 4 hours in vitro. After treatment, the lysates were immunoblotted under non-reducing conditions using the indicated antibodies. (E) The PC6 assays were performed in fractions from (4D) in the presence or absence of 200 μM NMN. Images or Blots are representative of three independent experiments. Data are presented as the means ± SD from three independent experiments. Statistical analysis was performed using Student’s t test.

**Figure S5.**
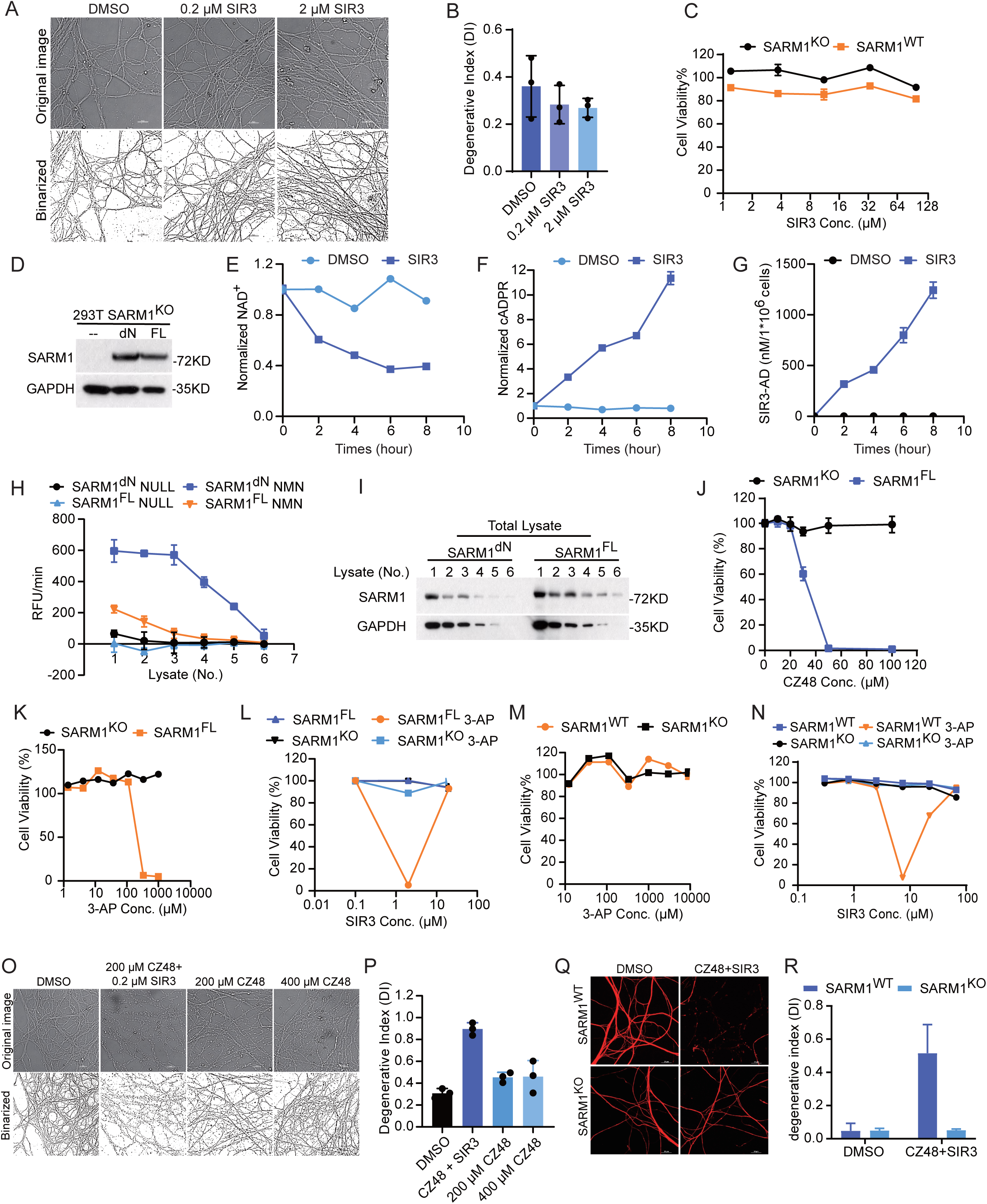
SIR3 induces sarmoptosis and axon degeneration through a two-step activation process, related to Figure 4. (A and B) Representative images of embryonic DRG neuron axons after treatment with different concentration of SIR3, as indicated, for 72 hours. (A) These binarized images are displayed to enhance the visualization of axonal integrity during the analysis of the degeneration index. (B) Quantification of DRG axonal degeneration. (C) 293T SARM1^-/-^ and WT cells were treated with different doses of SIR3 as indicated. Cell viability was measured with the CellTiter-Glo assay. (D) Western blot of the expression of dN-SARM1 and FL-SARM1 in HEK-293T SARM1^-/-^ cells. (E-G) 293T FL-SARM1 cells were treated with SIR3 (7 μM). Metabolites (E) NAD^+^, (F) cADPR, and (G) SIR3-ADPR were quantified by LC/MS at indicated times after treatment. (H and I) (H) The PC6 assay and (I) western blot analysis was employed to assess the SARM1 activity and the protein levels of SARM1, respectively, in lysates of different concentrations of 293T dN-SARM1 and 293T FL-SARM1. (J and K) 293T SARM1^−/−^ and FL-SARM1 cells were treated with varying concentrations of (J) CZ48, (K) 3-AP, as indicated. Cell viability was measured with the CellTiter-Glo assay. (L) 293T SARM1^−/−^ and FL-SARM1 cells were treated with 3-AP (100 μM) or a combination of 3-AP with varying concentration of SIR3 as indicated. Cell viability was measured with the CellTiter-Glo assay. (M) 293T SARM1^−/−^ and WT cells were treated with varying concentration of 3-AP as indicated. Cell viability was measured with the CellTiter-Glo assay. (N) 293T SARM1^−/−^ and WT cells were treated with 3-AP (900 μM) or a combination of 3-AP with varying concentration of SIR3 as indicated. Cell viability was measured with the CellTiter-Glo assay. (O and P) (O) Representative images of embryonic DRG neuron axons after treatment with different concentration of CZ48 or CZ48 + SIR3, as indicated, for 72 hours. These binarized images are displayed to enhance the visualization of axonal integrity during the analysis of the degeneration index. (P) Quantification of DRG axonal degeneration corresponding to (O). (Q and R) Representative images of axons from WT and SARM1 KO cultured DRG neurons at 80 hours after treatment with DMSO or CZ48 + SIR3. Immunofluorescent staining with antibodies against neurofilament (red). (R) Quantification of DRG axonal degeneration corresponding to (Q). Images or Blots are representative of three independent experiments. Data are presented as the means ± SD from three independent experiments. Statistical analysis was performed using Student’s t test.

**Figure S6.**
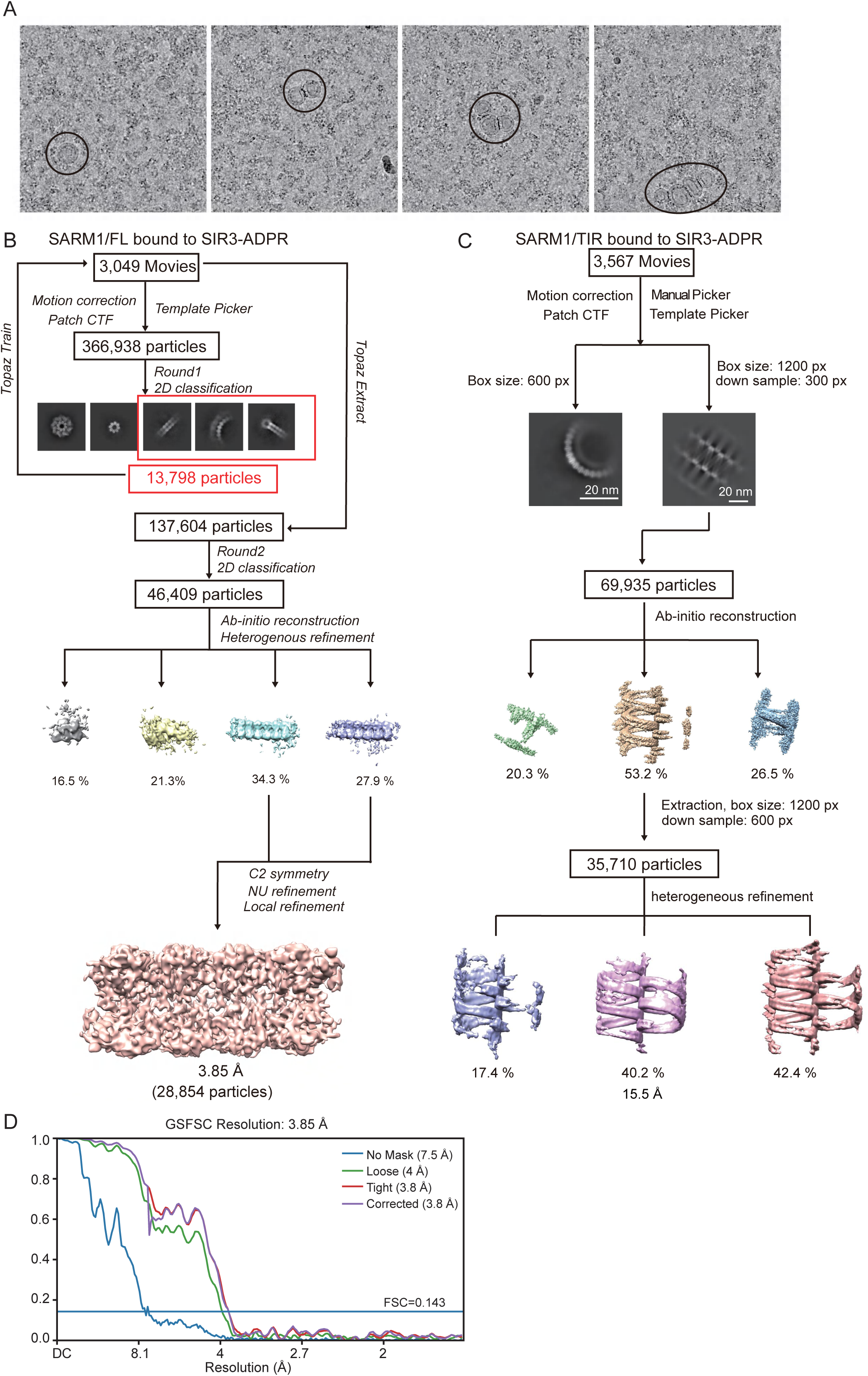
Cryo-EM structure determination, related to Figure 5. (A) Representative cryo-EM micrographs of SARM1FL incubated with SIR3-ADPR. (B) Cryo-EM workflow for SARM1 FL bound to SIR3-ADPR. 2D averages boxed in red are subjected to 3D reconstruction. (C) Cryo-EM workflow for SARM1 TIR domain bound to SIR3-ADPR. (D) Gold-standard FSC curves for the final map shown in (B).

**Figure S7.**
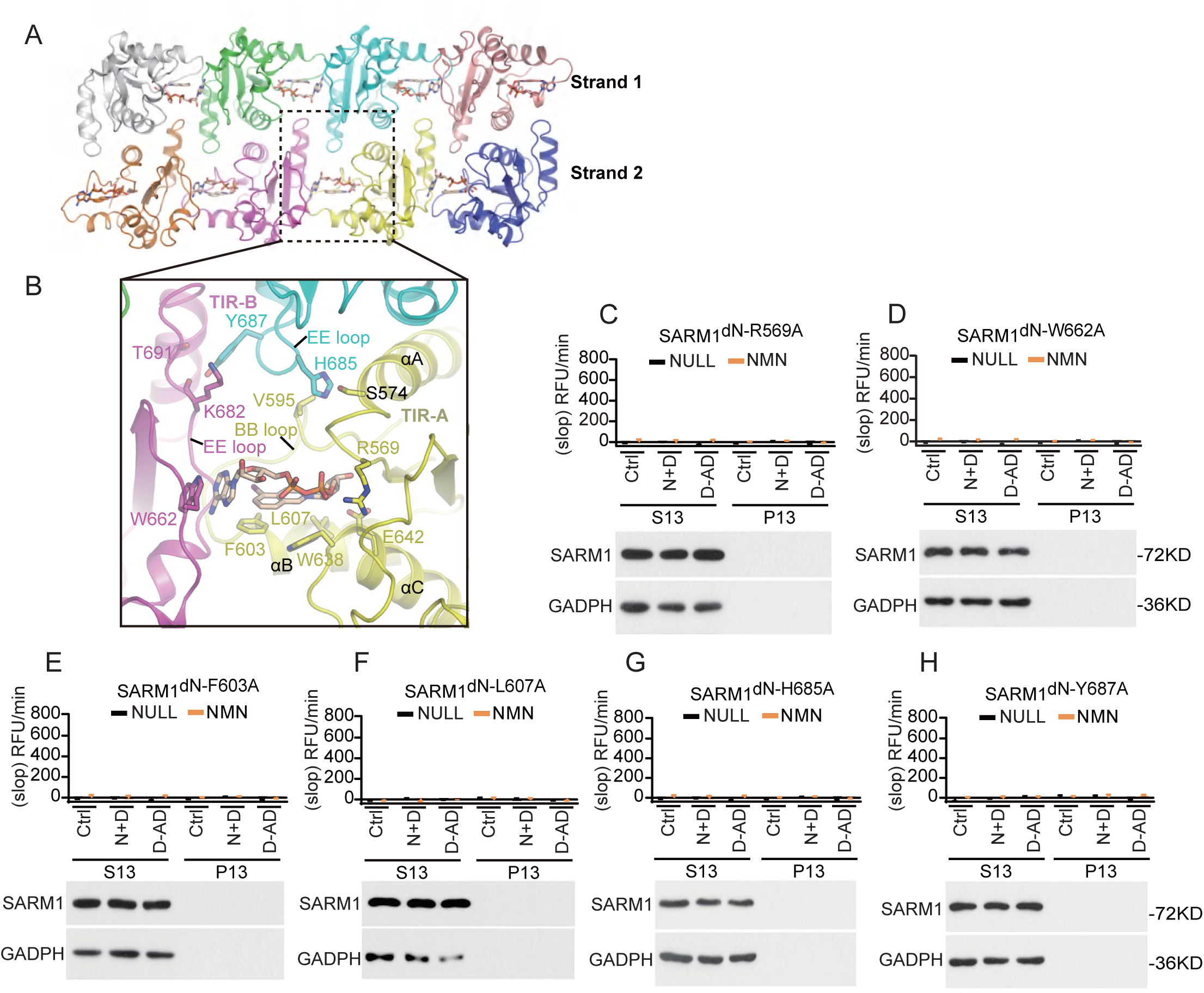
The formation of two antiparallel strands of the TIR domain of SARM1 is essential for SARM1 activity and consensates formation, related to Figure 5. (A and B) Detailed interactions between 1AD and the TIR domain of SARM1 (PDB: 7NAK). The EE loop is involved in interactions between two antiparallel strands of the TIR domain. (C-H) Effects of orthostatic-site and EE-loop mutations on hSARM1 NADase activity. SARM1 mutants’ lysates were treated with DMSO, SIR3 + NAD^+^, or SIR3-ADPR, followed by centrifugation, the S13 and P13 fractions were subjected to western blot and the PC6 assay in absence or presence of 200 μM NMN. Data are presented as the means ± SD, n = 3 independent experiments.

**Figure S8.**
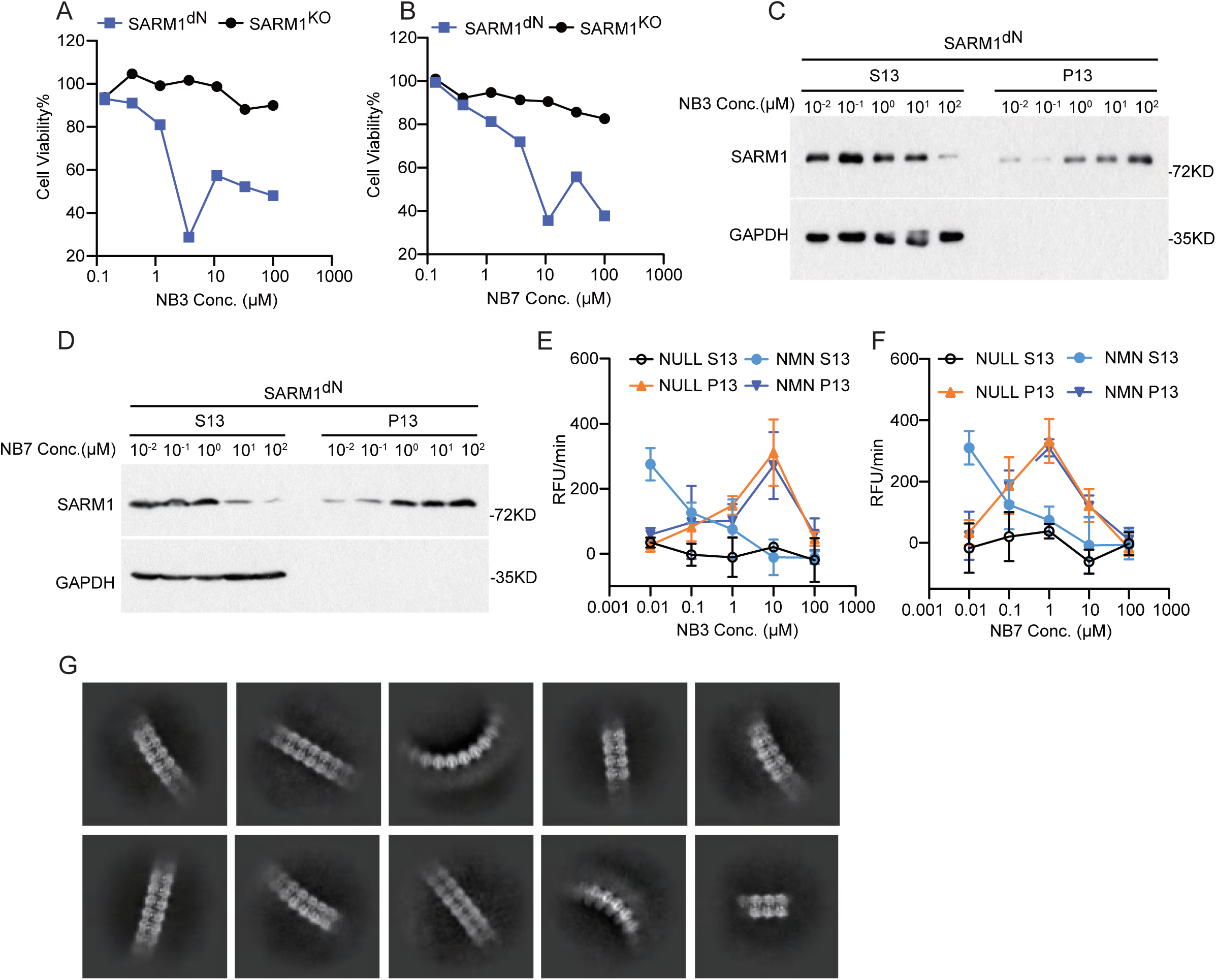
Reported SARM1-TIR inhibitors can also induce SIR3-like SARM1 activation, related to Figure 6. (A) SARM1^dN^-overexpressing HEK-293T cells were treated with varying concentration of NB-3 as indicated. Cell viability was measured with the CellTiter-Glo assay. (B and C) dN-SARM1-containing lysates were incubated with NAD^+^ (500 μM) and different doses of NB-3 as indicated, followed by centrifugation. After that, (B) all S13 and P13 were applied to western blots to assess SARM1 levels. (C) The PC6 assays were also performed in all S13 and P13 in the presence or absence of 200 μM NMN. (D) SARM1^dN^-overexpressing HEK-293T cells were treated with varying concentration of NB-7 as indicated. Cell viability was measured with the CellTiter-Glo assay. (E and F) dN-SARM1-containing lysates were incubated with NAD^+^ (500 μM) and different doses of NB-7 as indicated, followed by centrifugation. After that, (E) all S13 and P13 were applied to western blots to assess SARM1 levels. (F) The PC6 assays were also performed in all S13 and P13 in the presence or absence of 200 μM NMN. (G) Representative 2D class averages from cryo-EM data-sets of SARM1 incubated with DSRM-ADPR. Images or Blots are representative of three independent experiments. Data are presented as the means ± SD from three independent experiments.

## Notes

### Competing Interest Statement

Xiaodong Wang is a co-founder and consultant for Sironax Inc., a start-up company is developing therapeutic agents for neurodegenerative diseases.

### Summary of Updates

In order to improve the image resolution

